# Working memory expands shared task representations in cortex

**DOI:** 10.1101/2025.09.29.679345

**Authors:** E. Mika Diamanti, Lucas Pinto, Manuel Schottdorf, Abigail A. Russo, Stephan Y. Thiberge, Carlos D. Brody, David W. Tank

## Abstract

Cognition is thought to emerge from the flexible organization of neural activity, yet how this organization reconfigures across behaviors varying in cognitive load remains unclear. We investigated how the structure of working-memory representations in the cortex compares to task representations that do not involve working memory. We used a task-switching paradigm in virtual reality, where mice alternated between a navigation-based working-memory task and a simpler task with matched sensorimotor demands. During behavior, we simultaneously imaged three cortical areas: higher visual area AM, and two association areas—premotor (M2) and retrosplenial cortex. At the single-neuron level, trial-averaged activity appeared similar across tasks. However, pairwise correlations decreased during the working-memory task, particularly in association areas. In addition, the corresponding linear task subspace explained the variance of both tasks equally well, whereas the simpler task subspace failed to do so, suggesting an asymmetric relationship between them. Nonlinear dimensionality reduction revealed a shared low-dimensional structure across tasks. Yet, the organization of neuronal firing fields along this shared structure accounted for the difference in pairwise correlations: in the working-memory task, firing fields were more disjoint, especially among neurons in association areas that formed sequences along the memory dimension. Moreover, the degree of overlap between these firing fields predicted the mice’s behavioral reliance on working memory. We conclude that behaviors varying in cognitive demands are supported by a single low-dimensional neural structure, which can expand or contract depending on cognitive load. We thus provide a framework for how task representations across the cortex reconfigure to support cognitive processes.

## Introduction

Animals are capable of a remarkable range of behaviors. Strikingly, the coordinated neural activity that supports many of these behaviors often organizes into a low-dimensional structure^1–3^. Yet, how such structures relate across behaviors remains unclear, particularly in the context of cognition. Consider, for instance, working memory (WM), the short-term retention and manipulation of information^4^. We may or may not use WM as we make decisions or navigate the world. Does WM recruit a distinct neural structure, or does it reshape the existing one?

Numerous studies have provided substantial insight into the neural mechanisms underlying WM^5–12—^ from the sustained activity of individual neurons^13^ and their mixed selectivity^14^, to ‘activity silent’ synaptic mechanisms^15^, or other proposed population-level mechanisms, such as the sequential activity patterns^16,17^, attractor dynamics^18^, and the dynamic functional connectivity among neural ensembles^19^. However, it is not known how the neural structure of WM-dependent behaviors reconfigures when WM is absent.

At first sight, one might consider WM to be an additional task component, such that behaviors with and without WM would be associated with distinct neural representations. Another possibility is that WM is embedded in a high dimensional neural state space, as previously suggested in cortex^20^. A third possibility is that certain areas, such as the hippocampus, create low dimensional representations of WM variables^21^, forming conjoined cognitive maps^22^, of both spatial^23^ and more abstract task-relevant domains^24–27^. In that case, behaviors not requiring WM may rely on the same low dimensional representation, albeit compressed along the WM dimension.

We hypothesized that the principles underlying hippocampal cognitive maps of WM^21^, can extend to any brain area containing such representations, and examined how they reconfigure in the absence of WM. We focused on three cortical regions previously shown to belong to distinct functional clusters^28^: 1. higher visual area AM^29^; 2. retrosplenial cortex (RSC), involved in the encoding of visual stimuli^30–32^, spatial information^33–35^, context discrimination^36^, or history- and value-related signals^37^; and 3. premotor cortex (M2), associated with anticipatory activity^38^ and preparation of upcoming movements ^39^, timing ^40^, multisensory integration ^41^ and the encoding of information during the delay period of WM tasks^42,43^.

We studied WM in the context of evidence accumulation, which requires maintaining and updating partial information in memory until a decision is reached^44^. To isolate the effects of WM, we adopted a task-switching protocol in virtual reality^28^, where mice alternated between a navigation-based, evidence accumulation task and a similar navigation task that did not require WM. As in an earlier non-navigational paradigm^20^, trial-averaged choice-selective sequences were similar across tasks. However, in line with the widefield imaging results from our group^28^, pairwise correlations were significantly lower in the WM task than in the simpler one. Using non-linear dimensionality reduction, we formulated a geometric explanation for the reduced pairwise correlations and for how the same neural population can represent tasks with distinct cognitive demands. The decrease in correlations was reflected in the more disjoint organization of neuronal firing fields along the low-dimensional neural structure — the manifold firing fields. In parallel, the additional cognitive load was expressed as an expansion of the structure along the working-memory dimension.

Across mice, greater difference in the organization of manifold firing fields between tasks corresponded to increased behavioral reliance on evidence accumulation.

Taken together, our results demonstrate that tasks differing only in cognitive demands share a common low-dimensional structure that can expand along the WM dimension, depending on behavioral needs.

## Results

### Simultaneous cellular-resolution recordings in distinct cortical areas during tasks with different cognitive demands

We trained mice on a virtual reality task-switching protocol^28^, where they alternated between a WM task—the ‘towers’ task—and a simpler, visually guided task using the same environment (Figure 1a). In both tasks, mice navigated a 330 cm-long T-maze composed of three epochs: a 30-cm start, a 200-cm cue region with brief (200 ms) visual towers, and a 100-cm delay. In the towers task, mice accumulated evidence (the difference in tower count on the left versus the right side), retained this information and/or their decision during the delay, and turned toward the side with more cues to receive a 10% sucrose reward ^45^. By contrast, the visually guided task did not require WM because, in addition to the visual towers, a salient landmark visible from the start indicated the correct side throughout the trial. Only performance in the towers task was modulated by evidence, with higher accuracy on high evidence/ easy trials (Figure 1c, Suppl. Figure 1a). Crucially, motor variables such as the view angle or the turn onset were identical between the two tasks (Figure 1d, Suppl. Figure 1b-d), ensuring that any observed differences in neural activity were not driven by task-specific motor behavior.

**Figure 1:**
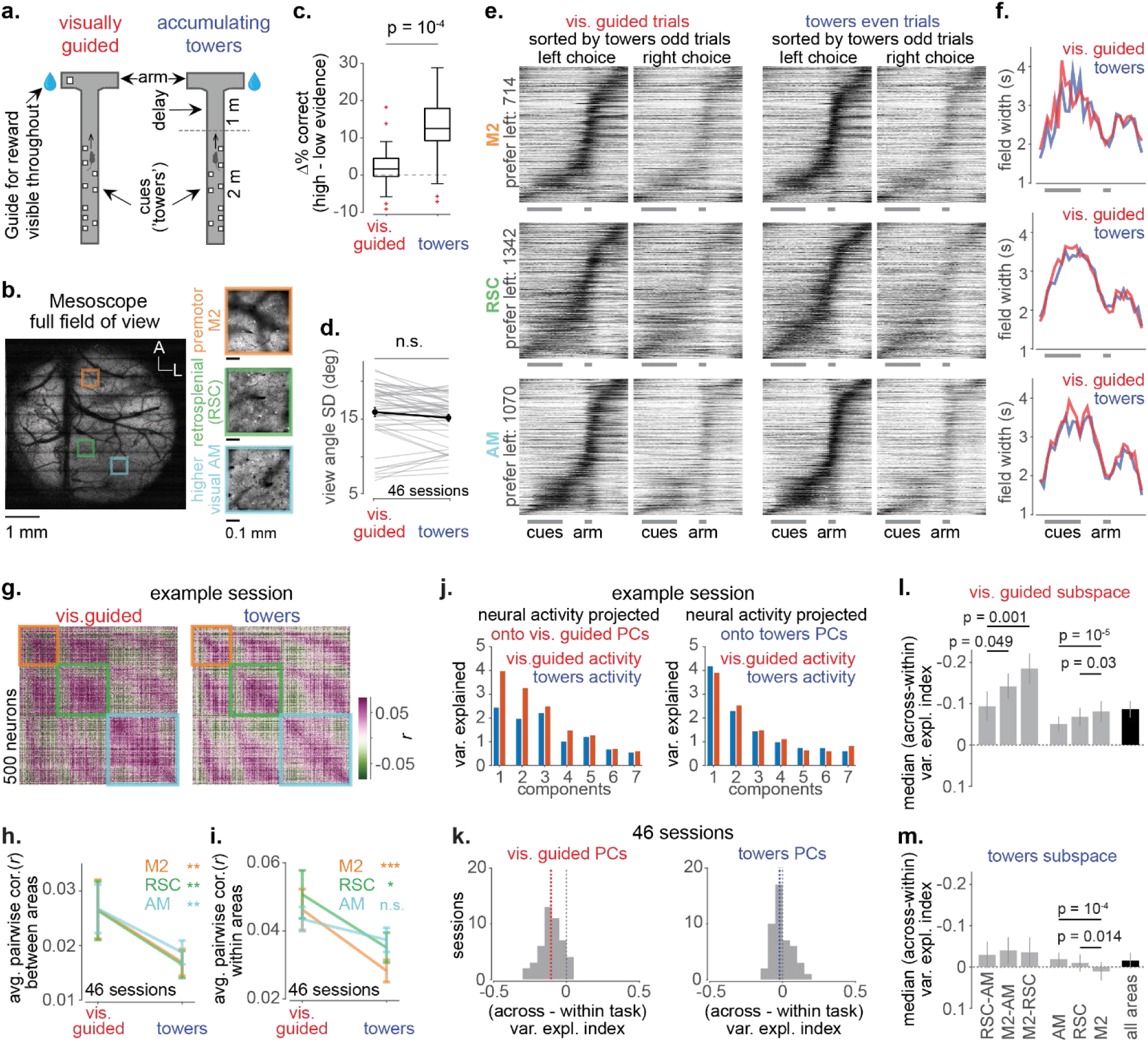
Choice-selective sequences across the cortex are the same in two tasks varying only in cognitive demands, but pairwise correlations differ. (a) Schematic of the task-switching behavioral protocol. Animals navigate a 330 cm T-maze, performing in blocks either a working memory task (right; ‘accumulating towers’) or a visual discrimination task (left; ‘visually guided’). (b) Mean image of a 6 mm craniotomy averaged across a z-stack of 20 imaging planes placed 40 μm apart. The image was acquired with a random-access two-photon microscope at full field of view (scalebar: 1mm; A: anterior; L: lateral). Superimposed squares denote the 500 µm x 500 µm imaging planes from one session, targeted on M2 (orange), RSC (green) and higher visual AM (cyan) Right: images of the imaging planes from the three areas averaged across 2,000 imaging frames (scalebar: 0.1 mm). (c) Distributions of the difference in performance in trials with high (Δ(#Left - #Right) cues > 5) versus low (Δ(#Left - #Right) cues ≤ 5) evidence in the two tasks, indicating the median (central mark), the 25^th^ (bottom edge) and the 75^th^ percentile (top edge). Whiskers extend to the most extreme datapoints that are not outliers. Red crosses: outliers. (p = 10^-4^, mixed effects model) (d) Trial-averaged view angle standard deviation (SD). Gray line: individual sessions (n = 46). Black line with filled circles: mean across sessions. Error bars: S.E.M. p-value from mixed effects model. (e) Cross-validated left choice-selective sequences in the visually guided task (left two columns) and the towers task (right two columns) in premotor M2 (top; 714 out of 1,899 neurons), RSC (middle; 1,342 out of 4,149 neurons) and AM (bottom; 1,070 out of 2,635 neurons). Neurons in both tasks were sorted based on the order obtained from the towers odd trials (‘train’ set). The plotted mean firing fields were obtained from all trials in the visually guided task and from the even-numbered trials in the towers task (‘test’ set). Sequencies span the whole trial plus the intertrial interval. x-axis gray scale bars: cues, arm. (f) Mean firing fields averaged across all neurons preferring a given position in the maze obtained from the test sets of each task (same area order and task color as in (e)). (g) Pairwise correlation matrix in the towers task (right) versus the visually guided task (left) from one example session, calculated across all possible pairs of recorded neurons (n = 500). Colored squares along the diagonal demarcate the within area pairwise correlations (orange: M2; green: RSC; cyan: AM). (h) Average across-area pairwise correlations in the towers versus the visually guided task for each area (p values from mixed effects model, M2: p = 0.005, RSC: p = 0.005, AM: p = 0.008; error bars indicate S.E.M.). (i) Average within-area pairwise correlations in the towers versus the visually guided task for each area (p values from mixed effects model, M2: p = 0.0002, RSC: p = 0.01, AM: n.s.; error bars indicate S.E.M.). (j) Towers even trial-to-trial neural activity variance (blue) or visually guided even trial-to-trial neural activity variance (red) explained by the first seven principal components of the visually guided task (left) or the towers task (right) from the same example session as in (g). (k) The variance explained index was defined as the normalized difference in the variance explained by a given task subspace when projecting data from the other task (‘across’) versus data from the same task (‘within’). The distribution across sessions of the variance explained index for the visually guided task was skewed towards negative values (left, median ± SEM: -0.11 ± 0.02; dashed red line indicates the median), unlike the distribution for the towers task which was centered around zero (right, median ± SEM: -0.02 ± 0.02; dashed blue line indicates the median). The two distributions are significantly different (p = 10^-10^, mixed effects model). (l) Median variance explained index of the visually guided subspace across sessions obtained from individual correlation submatrices, either within areas (M2, RSC or AM) or across areas (M2-RSC, M2-AM, RSC-AM). The total number of components used explained 20% of the variance in all cases. (p values are from mixed effects model). Error bars: S.E.M. (m) Same as in (l) for the towers task.

We recorded neural activity at cellular resolution from L2/3, using a random-access two-photon microscope with a 6 mm field of view^46^ (Figure 1b, Suppl. Figure 2). In nine transgenic mice expressing GCaMP6s, we targeted three cortical areas on the right hemisphere—premotor cortex (M2), retrosplenial cortex (RSC), and higher visual area AM— previously shown to form distinct functional clusters during this task ^28^. Across 46 sessions, we recorded 7,899 neurons in M2, 10,489 in RSC, and 8,630 in AM. Functional criteria confirmed accurate targeting of the posterior areas. Consistent with previous findings ^31^, ∼15% of neurons in AM and ∼5% in RSC were cue-locked. We also identified cue-locked neurons in M2, though their responses exhibited longer latencies (Suppl. Figure 3). Aside from these regional differences, cue-locked responses did not differ between the two tasks.

### Choice-selective sequences are similar in the two tasks

We next asked whether choice-selective sequences of sufficiently active neurons (with at least 0.1 transients/trial) differed between tasks (Figure 1e, Suppl. Figure 4a,b). For each neuron, we estimated firing fields separately for correct left and right trials using a cross-validated approach. Based on towers odd trials, neurons were classified as: (1) left-preferring, if peak activity on left trials was significantly higher (paired t-test); (2) right-preferring, in the opposite case; or (3) non-selective if no difference was found. Firing fields from towers even or all visually guided correct trials were then ordered by the peak positions obtained from towers odd trials. Consistent with previous work, neurons in all areas and tasks formed sequential activity spanning the maze and inter-trial interval, with RSC neurons tiling the trial more uniformly than those in M2 or AM^17^ (Suppl. Figure 4c). In addition, recordings from the right hemisphere yielded a slight bias toward left-preferring cells (M2: +6%, RSC: +4%, AM: +6%). However, choice-selective sequences did not differ between tasks: sequence patterns (Suppl. Figure 4c), field widths (Figure 1f), and peak firing positions (Suppl. Figure 4d) were preserved. Thus, trial-averaged activity of active neurons was nearly identical across tasks, indicating that the added WM demands in the towers task had no effect on choice-selective sequences.

### Pairwise correlations decrease in magnitude in the working memory task

Although mean firing fields were similar across tasks, task-related differences may be evident in the trial-by-trial correlations of neural activity. Prior work showed that more cognitively demanding tasks reduce noise correlations^47^. Consistent with this, large-scale pairwise correlations between cortical areas were lower in the WM (towers) task than in the visually guided task^28^. We therefore asked whether a similar reduction would be evident in the overall pairwise correlations at the single-neuron level.

For each session, we took the correct trials, matched their numbers between tasks, and computed Pearson correlation coefficients between single-trial neuronal traces across M2, RSC and AM. Correlations were visualized as square matrices, with diagonal submatrices representing within-area correlations and off-diagonal ones the between-area pairs (Figure 1g). Overall, pairwise correlations were lower in the towers task than in the visually guided task (Figure 1g, Suppl. Figure 5b), with both positive and negative correlations showing reduced magnitude (Suppl. Figure 5a, d, e). Despite this, correlation structure was similar between tasks, as indicated by a high R² across correlation matrices (Suppl. Figure 5c).

We next examined area-specific effects (Figure 1h, i). Consistent with our mesoscale wide-field data^28^, between-area correlations were significantly lower in all regions (M2: *p* = 0.005; RSC: *p* = 0.005; AM: *p* = 0.008; mixed-effects model). Within-area effects varied: M2 and RSC showed significant reductions in the towers task, whereas AM did not (M2: *p* = 0.0002; RSC: *p* = 0.01; AM: *p* = 0.1). These changes were not explained by differences in mean firing rates (Suppl. Figure 5f, g).

In sum, neuron pairs remained coactive across tasks, but the strength of their correlated activity decreased during the more cognitively demanding behavior.

### Only the working memory task subspace captures activity in both tasks

Given the lower pairwise correlations in the towers task compared to the visually guided task, we hypothesized that the neural subspaces defined by each task’s correlation matrix might also differ. We performed cross-validated eigendecomposition (equivalent to PCA on z-scored single-trial data) to estimate principal components (PCs) for each task. The first 15 PCs from odd trials captured ∼20% of the variance in each task, with no major differences except for PC1 (Suppl. Figure 6a).

To assess subspace similarity, we projected each task’s activity from the held-out, even trials onto each task’s subspace and estimated the variance captured by each individual PC (Figure 1j, example session – the first 7 PCs shown). Visually guided PCs better captured their own task’s activity than that of the towers task, whereas towers PCs explained both tasks equally well. We verified this by plotting the trial-to-trial activity of each task projected onto the first PC (Suppl Figure 6b left) or the second PC (Suppl Figure 6b right) of each task subspace. Along the towers axes, projected activity from both tasks spanned similar ranges (x-axes in Suppl. Figure 6b), but along the visually guided axes activity from the visually guided task was more dispersed than that from the towers task (y-axes in Suppl. Figure 6b).

To quantify this effect across sessions, we computed a variance explained (VE) index: the normalized difference between across-task and within-task variance explained by the first 15 PCs of each subspace^48^ (total number of PCs did not affect the result – Suppl. Figure 6c). A VE index of zero indicated that a subspace explained activity in both tasks; negative VE index indicated that a subspace better explained its own task. The distributions of VE indices differed significantly between task subspaces (Figure 1k, p = 10^-10^, mixed effects model). The towers-task subspace had a VE index near zero (median ± SEM: -0.02 ± 0.02), while the visually guided subspace had a negative VE index (median ± SEM: -0.11 ± 0.02). This finding suggests that only the WM-task subspace may be large enough to account for neural activity in both tasks.

### Task subspaces in association areas differ more than in higher visual

We next asked whether this asymmetry observed between task subspaces was present in the within-and the between-area pairwise correlations of individual areas (Figure 1l-m; Suppl. Figure 6d-i). We repeated the eigendecomposition on within-area correlations and applied SVD for between-area matrices. To compare across areas, we subsampled equal neuron counts 10 times and averaged VE indices across iterations. In the visually guided task, M2 had the most negative VE index among areas (vs. RSC: *p* = 0.03; vs. AM: *p* = 10⁻⁵), and between-area VE indices involving M2 were more negative than RSC–AM pairs (M2– RSC: *p* = 0.001; M2–AM: *p* = 0.049). In contrast, towers-task subspaces showed smaller VE index magnitudes across all areas, though M2 still differed significantly from AM (*p* = 10⁻⁴) and RSC (*p* = 0.014). No differences were observed in between-area VE indices for the towers task. Overall, subspace asymmetry was strongest in M2 and RSC, and in their pairwise interactions, suggesting that the WM subspace in these association areas may be broader than that of the simpler task, in contrast to AM.

### Population activity is organized in a low dimensional structure, with single-neuron firing fields overlapping less in the working memory task than in the simpler task

Next, we asked whether activity in both tasks could be captured by the same, potentially low-dimensional structure. Nevertheless, linear dimensionality reduction methods are inadequate for addressing this, because they require an exceedingly high number of dimensions to explain the variance in the data. More importantly, identifying the underlying shared structure between tasks most likely requires non-linear dimensionality reduction techniques^49^.

To investigate how population dynamics are organized trial-by-trial in each task, we applied MIND (manifold inference from neural dynamics^50^), a nonlinear method for reconstructing low-dimensional manifolds. For each cortical area, we extracted a low-dimensional manifold, and subsequently embedded it into an *n*-dimensional Euclidean space (*n* = 2–8 dimensions/latents, as in ^21^). We then reconstructed each neuron’s activity by mapping the latent dimensions back to neural state space and assessed reconstruction accuracy by correlating predicted and actual traces (Suppl. Figure 7a). With 5 dimensions, MIND achieved over 60% accuracy (Suppl. Figure 7b and ‘scores’ in Suppl. Figure 7c, M2: 0.60, RSC: 0.79, AM: 0.67). Although MIND is agnostic to the animal’s behavior, if the latent low-dimensional space indeed captures the task-relevant neural structure, task variables should vary smoothly along it. Indeed, both evidence and position formed smooth gradients along the first three latent dimensions across tasks and areas (Suppl. Figure 7c,d), similar to findings in hippocampus ^21^. In addition, similar to the mean behavioral fields (Figure 1), single-neuron activity tiled the latent space, forming ‘manifold firing fields’ (Figure 2b). Unlike behavioral firing fields though, the collective arrangement of manifold firing fields dictated how population activity evolves over time: activity follows smooth trajectories constrained by the manifold structure (Figure 2a).

**Figure 2:**
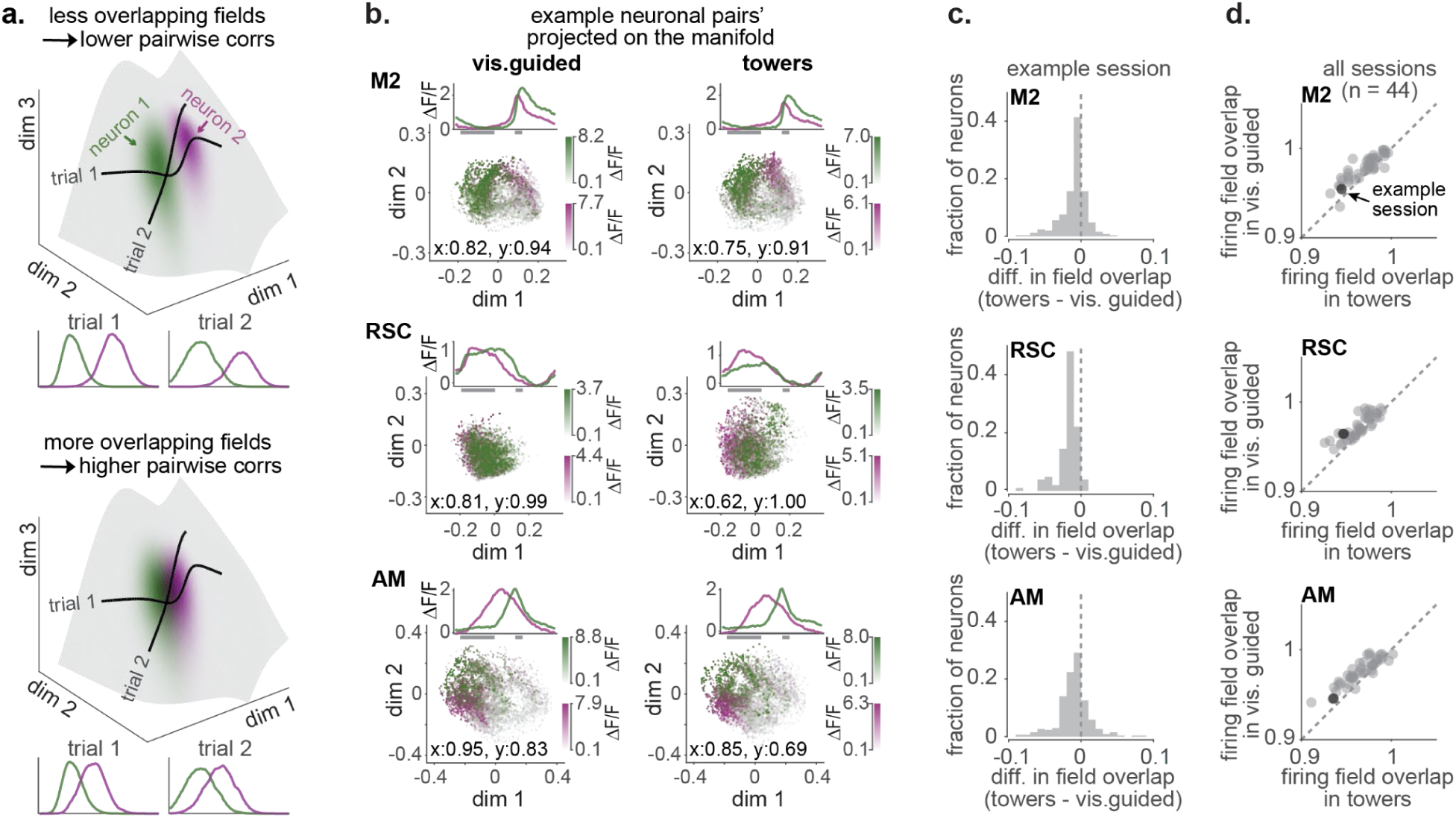
Single neurons form manifold fields that overlap less in the towers task compared to the simpler task. (a) Schematics showing how the arrangement of trial-to-trial activity on the manifold (‘manifold firing fields’) between neuron pairs can explain pairwise correlations, green: hypothetical manifold firing field 1, magenta: hypothetical manifold firing field 2. black lines: hypothetical single-trial trajectories of the network through the two neurons’ manifold firing fields (b) Example pairs of manifold firing fields and trial averaged fields (inset) in the towers (right) or the visually guided task (left), green vs magenta: example neuron 1 vs 2; x: overlap ratio along latent 1, y: overlap ratio along latent 2. Transparency of each cloud of points is weighed by the neuron’s ΔF/F magnitude (top: M2; middle: RSC; bottom: AM); gray bars under mean firing fields: cues and arm maze epochs **(c)** Normalized distribution of each neuron’s average difference in field overlap with all other neurons in the towers versus the visually guided task (top: M2, middle: RSC, bottom: AM). Same example session as in **(b)**. **(d)** Average firing field overlap in the towers against the visually guided task. Each data point corresponds to a single session (n = 44; black data point: same session as in **(b, c)**).

We focused on how single-neuron activity was organized on the manifold, reasoning that differences in pairwise correlations between tasks should be reflected in the overlap of their manifold firing fields: if two fields are adjacent but non-overlapping, the network trajectory can only pass through one at a time, leading to negative correlations. In contrast, fully overlapping fields imply that both neurons are active together or not at all, resulting in strong positive correlations (Figure 2a). We quantified the overlap between two manifold firing fields as the ratio of within-to between-field second moments for each neuron pair, where 1 indicates full overlap and ≪1 indicates minimal overlap (Suppl. Figure 7e). Most neuron pairs had greater overlap in the visually guided than the towers task (Figure 2b,c), a difference that was significant in all areas (Wilcoxon signed-rank test, *p* ≤ 10⁻¹⁷) and across all 44 sessions (mixed-effects model: M2 *p* = 10⁻⁵, RSC *p* = 10⁻⁴, AM *p* = 10⁻⁴; Figure 2d).

Therefore, although we identified a single lower-dimensional structure for both tasks, the organization of the manifold firing fields varied with task: they were more disjoint in the towers, reflecting lower correlated trial-to-trial variability.

### Neurons that tile the evidence space show the largest change in manifold field overlap between tasks

An earlier study demonstrated that hippocampal cognitive maps are instantiated as low-dimensional neural manifolds representing both physical (e.g., position) and abstract (e.g., evidence) variables in an orderly fashion^21^. Here, we show that the cortex also contains low-dimensional manifolds along which evidence varies smoothly. Notably, the tiling of these manifolds by individual neurons’ firing fields depends on the task. If the cortex indeed contains a cognitive map of abstract variables such as accumulated evidence—and given that evidence accumulation is the only factor distinguishing the two tasks — this leads to the hypothesis that the reduced overlap of firing fields in the towers task reflects the internal process of evidence accumulation. If so, the subpopulation of neurons encoding evidence should show the greatest task-dependent changes in firing field overlap.

To test this hypothesis, we first asked whether neurons in each area exhibited mean firing fields that collectively tiled the evidence space, forming sequences along the evidence axis (Fig. 3a). We focused on neurons active in the towers task (≥0.1 transients/trial) and computed trial-averaged activity across the evidence range [–12, 12] separately for odd and even trials. Neurons with positively correlated odd- and even-trial activity (p < 0.01; M2: 10%, RSC: 17%, AM: 17%) were selected and ordered by the evidence value at which their odd-trial activity peaked; this order was used to plot even-trial activity (Fig. 3a, right). All areas contained neurons tiling the evidence axis. We then repeated this analysis in the visually guided task, applying the same neuron set but sorting based on this task’s odd-trial activity. Sequences appeared noisier, indicating that these neurons encoded evidence more reliably in the towers task. We also estimated the mean field width of these neurons as a function of the evidence axis, by averaging across the ‘full width at half max’ (FWHM) of all even-trial averaged fields that peaked at a given evidence value. Field widths were generally larger in the towers task except at zero evidence (Fig. 3b), suggesting that these neurons tracked accumulated evidence, not just visual cues. Notably, the task difference in field width varied by area, with AM showing smaller differences than M2 and RSC (One-Way ANOVA with posthoc comparison, p-values: M2–RSC = 1; M2–AM = 0.02; RSC–AM = 0.02; Fig. 3c).

**Figure 3.**
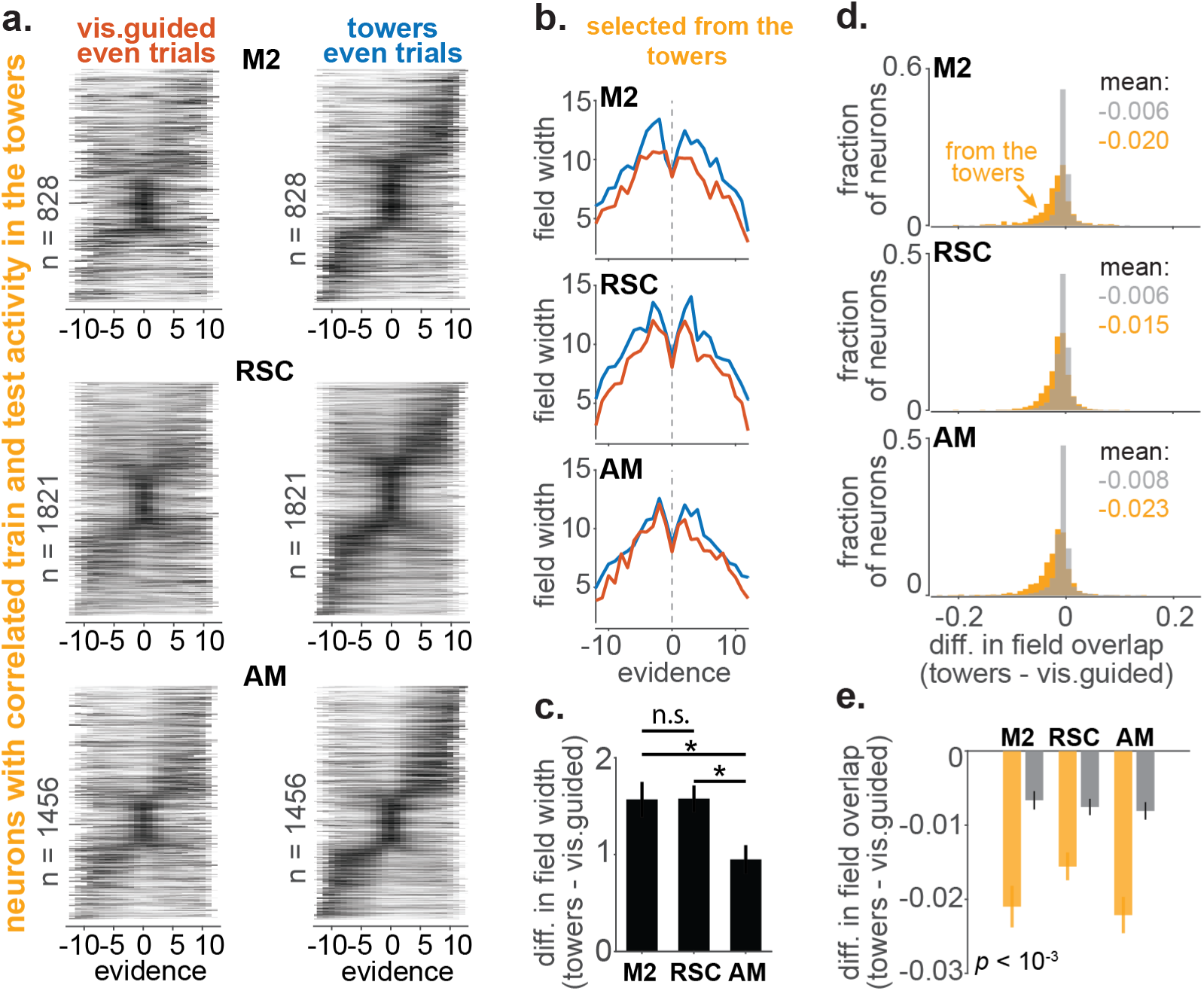
Neurons tiling the evidence space have firing fields on the manifold that overlap less in the working memory task than in the simpler task. **(a)** Cross-validated sequences as a function of the evidence axis obtained from neurons with correlated train and test activity in the towers task (top: M2, middle: RSC, bottom: AM). Plotted sequences are from the even trials of the towers (right) or the visually guided task (left), while the ordering of neurons was determined from the odd trials of each task. **(b)** Mean field width of the neurons in **(a)** as a function of the evidence axis in the towers (blue) or the visually guided task (red). **(c)** Difference in field width in the towers versus the visually guided task averaged across all evidence values. (1 star: p < 0.05, p value from one-way ANOVA with posthoc comparison; error bar: S.E.M.) **(d)** Normalized distribution of the average difference in manifold field overlap within the same manifold, for neurons participating in the evidence sequences in **(a)** (orange) or for all other neurons (gray), concatenated across sessions (n = 44, top: M2, middle: RSC, bottom: AM). **(e)** Mean difference in manifold field overlap in the towers versus the visually guided task averaged across sessions, for neurons participating in the evidence sequences in **(a)** (orange) versus all other neurons (gray) (p < 10^-3^ mixed effects model in each area; error bar: S.E.M).

The difference in firing field width was not due to selecting neurons based only on towers-task activity. When we repeated the analysis using neurons selected in the visually guided task (Suppl. Fig. 8a), selection rates were similar (M2: 9%, RSC: 13%, AM: 13%). As expected, these neurons showed less noisy evidence sequences in the visually guided task compared to the towers. However, unlike neurons selected in the towers task, their field widths were at most equal across tasks (Suppl. Fig. 8b), and the difference across areas was abolished (Suppl. Fig. 8c; One-Way ANOVA, *p* value: n.s.). These findings suggest that during the towers task, especially in M2 and RSC, a specific subset of neurons formed wider firing fields that tiled the evidence space.

We next examined how the subpopulation encoding evidence in the towers task was organized within the previously obtained low-dimensional neural structure. We compared changes in manifold field overlap for these neurons versus others from the same session (Figure 3d). Neurons participating in the towers evidence sequence were distributed towards more negative values, indicating larger overlap changes between tasks. In contrast, the distribution of other neurons was centered around zero, indicating smaller change. This was confirmed by comparing mean field overlap across sessions (Figure 3e; towers evidence neurons vs. others, mean ± S.E.M.: M2: -0.021 ± 0.003 vs. -0.007 ± 0.001; RSC: -0.016 ± 0.002 vs. -0.007 ± 0.001; AM: -0.022 ± 0.002 vs. -0.008 ± 0.001; mixed effects model; all p < 10⁻³). In addition, the difference in firing-field overlap between tasks was clearly evident along the axis that best encoded evidence within the low-dimensional latent structure (Suppl. Figure 8d; mixed effects model; all p < 10⁻³). Therefore, evidence is represented on the neural structure primarily through neurons that collectively encode evidence in the towers task. These neurons exhibit wider trial-averaged fields along the behavioral evidence axis and greater separation of manifold firing fields on a trial-by-trial basis.

### In association areas, working memory expands an otherwise shared structure underlying both tasks

And so, how similar is the structure between the two tasks? To answer this question, we constructed a separate manifold for each task and applied an alignment method previously used for aligning manifolds across subjects ^21,51^(Figure 4a). In essence, a decoder obtained from the latents of one task was tested on the same or the other task. But since in an n-dimensional space there are 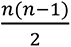 possible rotations that can be performed on an object while preserving its structure, we aligned each task latent space by choosing the rotation transformation that maximized decoding accuracy for position and evidence during the cues and the delay. We focused on 5-dimensional embedding spaces because they nearly maximized reconstruction scores (Suppl. Figure 7f). Alignment was done in both directions—using either the visually guided space as the reference frame while rotating the towers, and vice versa. If the two task structures are the same, decoders should yield similar results regardless of reference frame.

**Figure 4:**
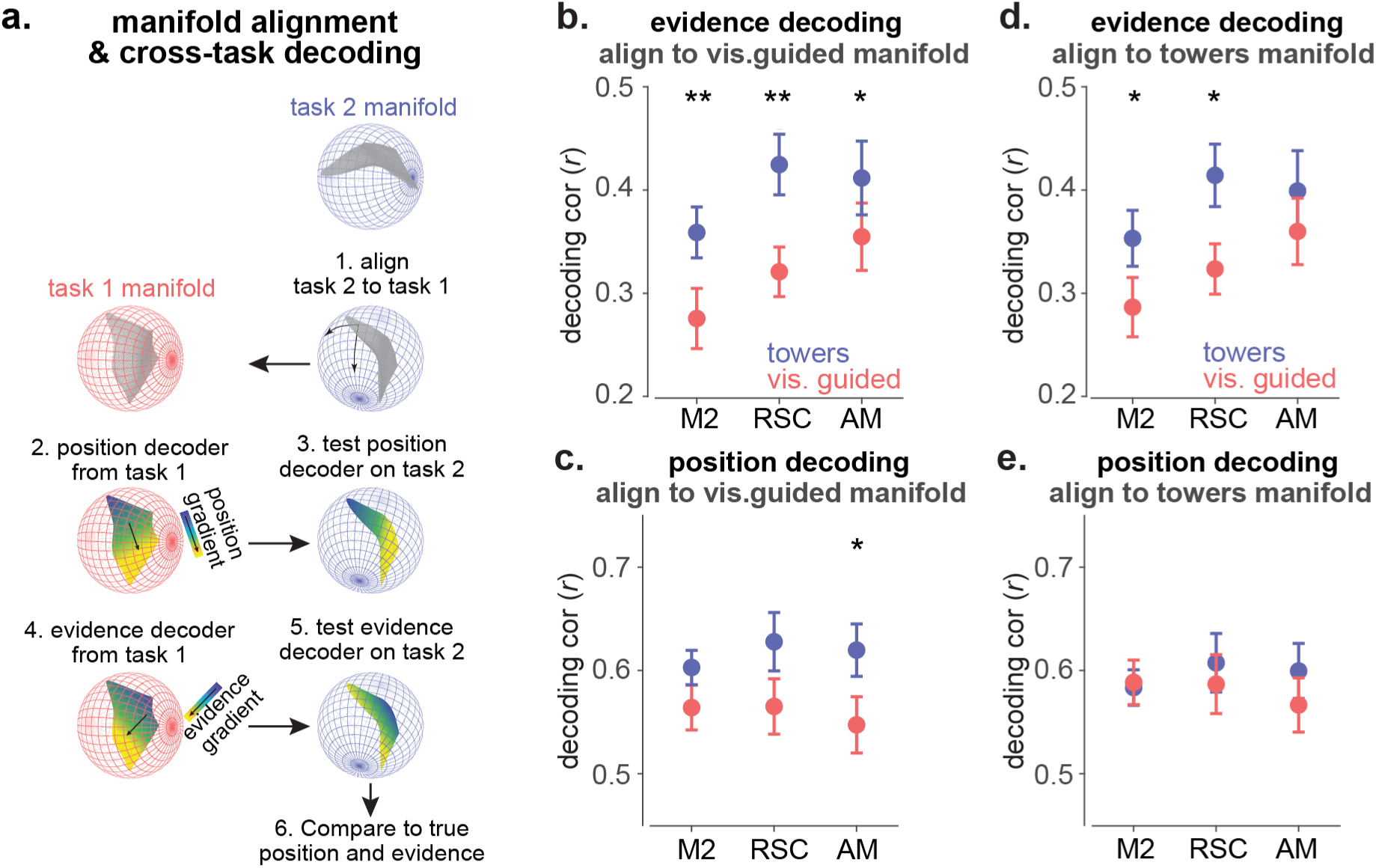
In association areas M2 and RSC, the towers task and the visually guided task manifolds have similar structure, but they vary specifically along the evidence gradient. **(a)** Schematics of the manifold alignment and cross-task decoding. The manifold from task 1 is used to train decoders of evidence and position. The decoders are then tested on held out trials from the same task or on trials from task 2, after aligning the task 2 manifold according to the behavioral gradients in task 1. **(b, c)** Alignment of the towers task manifold to the visually guided manifold. **(b)** Evidence decoding from the visually guided latents (red) or the rotated tower latents (blue). Evidence can be decoded better from the rotated towers latents than the visually guided latents (p value: M2: 0.006, RSC: 0.007, AM: 0.044, mixed effects model). **(c)** Same as in **(b)** for position. Position can be decoded better from the rotated towers latents than the visually guided latents only in AM (p value: M2: n.s., RSC: n.s., AM: 0.015, mixed effects model). **(d, e)** Alignment of the visually guided task manifold to the towers manifold. **(d)** Evidence decoding from the towers latents (blue) or the rotated visually guided latents (red). Evidence can be decoded better from the towers latents than the rotated visually guided latents in M2 and RSC (p value: M2: 0.027, RSC: 0.021, AM: n.s. , mixed effects model). **(e)** Same as in **(d)** for position. Position decoding accuracy is higher in the towers than the visually guided task only in AM (p value: M2: n.s., RSC: n.s., AM: 0.015, mixed effects model).

We first focused on evidence decoding by aligning the towers to the visually guided task (Figure 4b). Across areas, evidence was decoded more accurately from the rotated towers latents than the visually guided latents (*p* value: M2: 0.006, RSC: 0.007, AM: 0.044, mixed effects model). Similar results held when aligning the visually guided task to the towers task, particularly in M2 and RSC (Figure 4d, difference in decoding accuracy for each area, *p* value: M2: 0.027, RSC: 0.021, AM: n.s., mixed effects model). We then turned to the position decoders (Figure 4 c, e). Training on the visually guided latents yielded similar accuracy across tasks in M2 and RSC, but not AM (Figure 4c; *p* value: M2: n.s., RSC: n.s., AM: 0.015, mixed effects model). Training on the towers latents, however, led to matched decoding accuracy between tasks in all areas (Figure 4e; all *p* values: n.s., mixed effects model).

Overall, aligning one task or the other had no impact on decoding accuracy in M2 or RSC. Decoding accuracy for position was the same between the training and the aligned manifold (panels c. and e.), while evidence decoding was always better from the towers latents (panels b. and d.). In contrast, decoding in AM depended on the reference frame. Aligning the visually guided task to the towers yielded similar decoding accuracy between tasks for both position and evidence (panels d. and e.). But aligning the towers to the visually guided yielded higher accuracy for the towers task (panels b. and c.). We reasoned that the lower position decoding accuracy in AM’s visually guided reference frame (panel c) reflects variability in visual responses, which may be on or off depending on the guide’s position on a given trial. Hence, we could not dissociate evidence from position encoding in AM. By contrast, in M2 and RSC, evidence—but not position—was better decoded during the towers task, regardless of reference frame. This indicates a shared task structure in M2 and RSC. However, the fact that evidence is consistently better encoded by the towers latents implies that the corresponding population trajectories span a more expanded region of latent space than those in the visually guided task.

### The degree of overlap between task representations reflects how strongly behavior is influenced by evidence

So far, our data suggest that the towers task shares a similar representational structure with the visually guided task, but occupies a more expanded region along the evidence dimension. If so, then in mice whose behavior is less modulated by evidence—and thus more similar across tasks—the task representations should become even more alike. In other words, the manifold field overlap difference (Figure 2), and the linear subspace asymmetry (Figure 1) should all diminish. For each animal, we quantified how evidence accumulation influenced their behavior using the approach in Figure 1b and used this as a measure of behavioral similarity between tasks (Figure 5a). For example, Mouse 1 showed distinct behaviors across tasks (p = 0.008 two-sided Wilcoxon rank sum test), mouse 3 the most similar (n.s. two-sided Wilcoxon rank sum test), and mouse 2 intermediate (p = 0.015 two-sided Wilcoxon rank sum test). We defined the behavioral difference in evidence modulation as the difference between the 25th percentile of the towers distribution minus the 75th percentile of the visually guided distribution. We then plotted this ‘evidence modulation difference’ across mice against the VE index of the visually guided subspace (Figure 5b) and found a strong anticorrelation (*p* = 0.001, bootstrapping). As behavioral similarity increased, the asymmetry between linear task subspaces decreased. In contrast, the towers subspace explained both tasks well, regardless of the inter-subject behavioral variability (Suppl. Figure 9; *n.s.*). We then examined the relationship between behavioral similarity and the non-linear manifolds (Figure 5c). The difference in manifold field overlap between tasks (along the first latent) decreased with behavioral similarity (Figure 5c), reaching near zero for mouse 3 (*p* = 0.008, bootstrapping).

**Figure 5:**
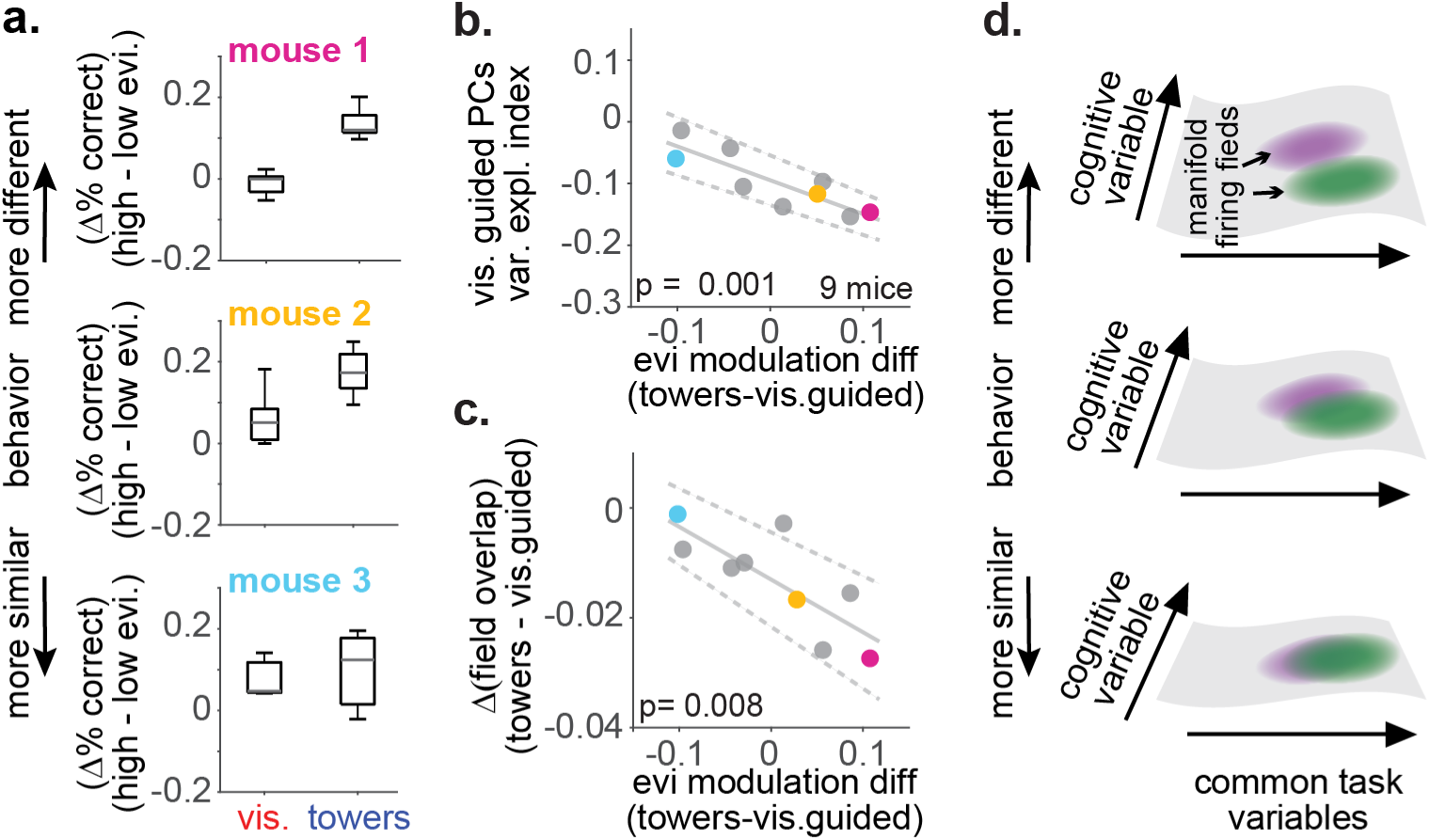
(a) Distributions of the difference in performance in trials with high versus low evidence in the two tasks (as in Figure 1c) for 3 example subjects. Subjects are ordered based on the difference between the bottom edge of the towers boxplot minus the top edge of the visually guided boxplot (‘evidence modulation difference’) **(b)** For each subject (n = 9), the evidence modulation difference (x axis) is plotted against the session-averaged variance explained index of the visually guided linear subspace (defined as in Figure 1k). The more different the behavior between tasks, the more negative the variance explained index (p = 0.001, from bootstrapping, solid line: fitted line averaged across bootstrapped iterations, dashed lines: ± 2 standard deviations). Colored datapoints: example mice shown in **(a),** gray datapoints: all other mice **(c)** For each mouse, the same x-axis as in **(b)** is plotted against the session-averaged difference in manifold field overlap in the towers versus the visually guided task along the first latent dimension (defined as in Figure 2). The more different the behavior between tasks, the smaller the overlap between firing fields on the towers manifold compared to the visually guided manifold (p = 0.008, from bootstrapping, solid gray line: mean slope and dashed gray lines: 2 SDs, from 1,000 bootstrapped iterations). **(d)** Schematics summarizing the effect of cognitive processes on the task representation shared between tasks with distinct cognitive demands.

In sum, behavioral variability across subjects is reflected in how well the visually guided subspace captures the towers task, and also in the degree of overlap between manifold firing fields. Particularly in the non-linear regime, stronger behavioral modulation by evidence, leads to more disjoint manifold firing fields (figure 5d). These results show that the neural representation capturing the ‘backbone’, i.e. the task variables common in both tasks, can expand or contract depending on the effect of abstract cognitive variables such as working memory.

## Discussion

We have shown that tasks with matched sensorimotor configuration, but distinct cognitive demands share a common low-dimensional neural structure. WM influences this structure by making manifold firing fields more disjoint. Together with the observed asymmetry in the linear domain—where the complex task explained the simpler one but not vice versa—these findings suggest that cognitive variables expand the low-dimensional task subspace. This expansion is also reflected in the widening of the behavioral fields along the evidence axis during WM, analogous to the widening of place fields along the expanded spatial dimension^52^. Even when analyzed separately, both tasks encoded information similarly, though sensory evidence was more strongly represented in M2 and RSC during the WM task. Finally, the degree of similarity between subspaces—captured in the linear domain by their ability to explain the other task, and in the nonlinear domain by the overlap of manifold firing fields— correlated with how similar the behaviors were across tasks.

Our cellular resolution data precisely replicates our earlier widefield study^28^ and extends it by providing a conceptual framework for the decrease in the magnitude of pairwise correlations during the WM task, while preserving their structure. As in the widefield imaging data by Pinto et al., pairwise correlations in the posterior cortical area AM did not change in magnitude between tasks. In addition, using cross-task, comparative analyses, we were able to identify differences between areas, despite the similarity in activity patterns at the single-neuron level. AM’s neural dynamics exhibited minor differences between tasks, suggesting that, unlike M2 and RSC, AM primarily encodes the low-level, common variables between tasks. This finding agrees with the optogenetic perturbations of the same study showing that performance was affected in both tasks when inactivating AM, but only during the WM task when inactivating M2 and RSC, and it further indicates that M2 and RSC, but not AM, play a role in WM. Thus, despite the widespread involvement of cortical areas during complex behaviors, we provide evidence for the functional segregation between areas of the dorsal cortex.

On the other hand, the contribution of both M2 and RSC to behaviors requiring WM is consistent with the idea of functional redundancy across distributed cortical networks^53^, which, similarly to motor planning^54^ may underlie the organization of cortical networks involved in WM. Within this distributed network, interareal interactions may play a prominent role in sustaining representations of WM ^20^. Our analysis of the between-area pairwise correlations and their associated linear subspaces point in the same direction. Future studies using more sophisticated models to infer interareal communication^55^ will provide further insight into the role of interareal interactions ^56^ within distributed cortical networks.

Our results also provide an explanation for the findings reported by other groups, such as the effects of prior cognitive experience on neural dynamics^47^ and the high dimensional linear embedding of WM^20^. Previous experience in a complex task requiring short-term memory recruits higher level cortical areas and decreases their pairwise correlations relative to subjects trained in a simple task only^47^. We propose that with prior cognitive experience the brain constructs a representation of the more complex task that it later reuses to represent the simpler but similar task. Similarly, task switching behavioral paradigms in which the two tasks are not in conflict, may recruit activity that would otherwise be absent if subjects were trained in one task only, by organizing neural dynamics in a single neural representation that reconfigures specifically along the dimensions representing cognitive variables. Such organization can explain the seemingly indistinguishable patterns of activity in the linear regime and the attribution of increasingly high dimensions to WM^20^. Nevertheless, using non-linear dimensionality reduction approaches we were able to extract low-dimensional embeddings that encompassed cognitive variables, and even detect a strong correlation between the organization of manifold firing fields along the very first latent dimension and the level of behavioral similarity between tasks.

Although originally coined for hippocampus^57^, task-dependent cognitive maps may provide a general framework for characterizing the influence of cognitive variables on neural dynamics. Here we show that similar principles of hippocampal computation^21^ also apply to certain areas of the dorsal cortex and in addition we propose a mechanism by which these areas can represent tasks with similar sensorimotor structure. Future studies should focus on the origins of such representations and how they evolve with learning.

## Methods

### Animals

All procedures were approved by the Institutional Animal Care and Use Committee at Princeton University under protocol 1910 and were performed in accordance with the Guide for the Care and Use of Laboratory Animals. We used 11 transgenic mice aged 10 to 16 weeks, 6 females and 5 males, from two transgenic lines that express the Calcium indicator GCaMP6s in cortical excitatory neurons (5 double transgenics: CaMKII-tTA; TRE-GCaMP6s line G6s2 – Jax stock # 007004 and 024742, and 6 single transgenics: GP4.3 – Jax stock # 024275). Mice were housed in groups of 2 to 6 and kept in a reversed light-dark cycle.

### Surgeries

Mice underwent sterile stereotaxic surgery to implant a 6 mm optical cranial window and a custom stainless steel headplate (∼1 g) under isoflurane anaesthesia (3% for induction, 1.5% for maintenance). Prior to any procedure, the mouse received 1 IP injection with 1 mg/kg meloxicam for analgesia, 2 mg/kg dexamethasone as anti-inflammatory, and with 0.5 ml of warm (37⁰C) saline to prevent dehydration. The animal was then placed into the stereotax, with its body lying on a heating pad, the animal’s eyes were protected with veterinary ophthalmic ointment and its temperature was monitored throughout the procedure with a rectal temperature probe. The fur over the surgical site was shaved and then scrubbed thrice with iodine followed by 70% ethanol. Using a pair of curved-tip fine scissors a ∼8 mm incision was performed in the AP direction over the midline and the skin was gently protracted until part of the lateral and posterior muscles and frontal bone were exposed (3-4 mm anterior of bregma). The periosteum was removed, and excess skin was glued at the edges of the exposed skull using tissue adhesive (3M Vetbond). To ensure that the head was perfectly flat relative to the stereotax surface, we made small adjustments in the pitch and roll of the head until lambda and bregma came into focus under a 40X centering scope with a cross-line reticle (KOPF model 1915). We used the same instrument to measure the distance between lambda and bregma and later the distance between bregma and the anterior and medial edge of the craniotomy. These measurements were subsequently used to register the craniotomy to the Allen CCF (see **Registration to the Allen Common Coordinate Framework**). A 6 mm craniotomy was performed over the right hemisphere. A 6 mm sterile disposable biopsy punch (Integra Miltex) was used to demarcate the edges of the craniotomy, with the anterior edge from bregma at ∼2.5 mm and the lateral edge at ∼ 4 mm from the midline. We then used a dental drill to drill over the craniotomy edges, and to loosen the sutures, especially those along the midline. Eventually, the bone was divided into 4 flaps bordered by the interfrontal, coronal and sagittal sutures, and removed one by one using curved fine forceps and an angled micro point. We next implanted the cranial window which consisted of 3 6-mm diameter round #0 (∼100 μm thick) glass coverslips glued together and bonded to a stainless-steel ring (OD: 6.5 mm, ID = 5.8 mm, thickness: 0.5 mm) using a UV-curing, clear, optical adhesive. An optical pole with magnets on one side was connected to the stereotax and was used to precisely position the window implant over the craniotomy and as parallel as possible to the head. The window implant was attached to the bone with dental cement (Metabond, Parkell). Finally, a second post with a headplate holder was connected to the stereotax and was used to position a stainless steel headplate featuring a curved bottom that fitted the shape of the skull. Thus, by aligning the head relative to the stereotax and precisely positioning the coverslip and headplate, we ensured that the headplate was flat and parallel to the coverslip and the surface of the brain, which is critical when imaging large fields of view.

### Two-photon, cellular-resolution imaging of 3 cortical areas simultaneously

Imaging over a large FOV was performed using a random-access two-photon microscope^46^. Briefly, the microscope has 2 main features: the scanning system consists of one resonant scanner and a pair of galvo scanners that allows the displacement of the fast scan line over the large FOV; a remote-focus unit which compensates for aberrations produced by the imaging objective (NA, 0.6, focal length 21 mm), and also enables fast focusing along the z axis, which would otherwise be impractical due to the bulky imaging objective. Two-photon illumination was performed with a Ti:Sapphire laser (Chameleon, Coherent) operating at 940 nm. The maximum power under the objective was 89 mW. Laser power was modulated with a Pockels cell (Conoptics). Depending on the mouse line and expression levels, laser power was set within the range of 40% and 80%. Imaging settings were controlled with ScanImage (SI; Vidrio Technologies). The imaging parameters and online motion correction were configured using an SI .usr and .cfg file and a custom Matlab function.

Although extra care was taken during surgeries to place the headplate and coverslip parallel to the craniotomy, small adjustments in the pitch and roll, of the order of 1 to 3 degrees, were still required to ensure that the cranial window was parallel to the objective. Before starting to image from a given mouse, we used a 1P, wide-field imaging path to focus on the surface of the brain. We then switched to the ‘alignment’ path. This path imaged the reflection of a red LED, which, after passing through a pinhole, was reflected from the cranial window onto the camera. A predefined, yellow outline on the image indicated at which position the reflection came from a surface parallel to the objective. The reflection from the cranial window was moved within the outline, by adjusting either the pitch of the headholder with a motorized linear stage (X-MCB2, Zaber) or the roll of the microscope’s objective using the microscope’s motors (thorlabs). The pitch, roll, as well as the height of the head holder, which was also adjusted using the motorized stage (19 -20 mm between the mouse’s paws and the headplate), were saved and retrieved in subsequent sessions.

On a typical imaging experiment, we first set the z-position and the pitch of the head holder to the previously saved values for that specific mouse. Next, we head-fixed the mouse and attached a 3-D printed light shield with a cut black balloon on top, onto the headplate using silicone elastomer (kwik-cast). Finally, we covered the cranial window with ultrasound gel (Sonigel, Mettler Electronics), which served as the immersion medium for the objective. We switched to the 1p pathway by sliding the dichroic mirror right above the objective and turned on a green LED for wide-field imaging. We used the motors of the microscope to rotate and lower the objective until the surface of the brain came into focus. We also adjusted the X and Y directions, until the wide-field FOV matched those from previous sessions. We switched to the 2P pathway by sliding the dichroic back in place. We then turned on the voice coil that carried the mirror of the remote focus unit, checked the beam power modulation and the fastZ actuator tuning with SI. We turned the photomultiplier tubes on and started imaging in full FOV mode, so that the whole 6 mm craniotomy would be visible under the 2-photon mesoscope. Imaging planes were typically 180 μm and 350 µm below the brain surface. To achieve maximum signal to noise ratio, we used the z-motors together with the remote focus, so that the imaging plane was lowered to a remote focus position that was close (∼50 um) - but not on - the focal plane of the remote focus mirror. With ROI mode enabled, we positioned 3 square-ROIs of 500 x 500 µm each, over the 3 areas of interest, based on the registration of the full FOV image to the Allen Common Coordinate Framework. We then ran a custom Matlab function that set up the online motion correction using the SI MotionManager, and configured SI for data acquisition. For the online motion correction, for each imaging ROI, SI acquired 10 zstacks of 41 planes each, spaced by 1 µm and centered around the imaging plane. The 10 volumes were first registered and then averaged, producing the final template for online motion correction. To track and correct the imaging plane’s z coordinate, we used SI’s MotionManager classes, MariusMotionEstimator and MariusMotionCorrector. Correction tolerance was set to 1 µm. Finally, just before starting to acquire, we switched on the two magnetic bases that provided extra stability for the motor stage. We acquired single planes from each FOV (n = 3) at a frame rate of 14.3 fr/s per plane.

### Registration to the Allen Common Coordinate Framework (CCF)

Mesoscope full FOV images were registered to the flattened Allen Brain Atlas API Mouse Brain (ccv3) using rotations and scaling. For each mouse, we marked bregma, B, and a second point, P, along the midline of the full FOV. With these measurements we estimated the rotation angle of the Atlas: 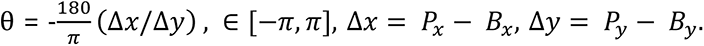 We then measured the distance from bregma to the anterior and lateral border of the craniotomy in pixels and converted it to mm using the pixels per mm ratio saved in the tiff header. These numbers were compared to the previously acquired measurements during surgery, and bregma was adjusted accordingly if numbers did not match. We then mapped the distance from bregma along the anterior-posterior and lateral-medial axis onto the Allen Brain image using the map’s pixels per mm ratio and cropped the map on the edges. Finally, the cropped map was scaled to the pixel size of the full FOV. We did not attempt to register using retinotopic maps, because within the 6 mm FOV that contained M2 only the anterior tip of primary visual cortex (V1) was usually exposed. Registration to the CCF was performed for any given mouse before the start of any imaging experiments during behavior. The registered full FOV was used to position the imaged ROIs on the right hemisphere over M2, RSC and visual AM.

### Virtual reality setup

The VR setup was similar to the one described previously^45^. Briefly, it consisted of a custom 3d printed cup, an optical flow sensor, a DLP projector, and a toroidal screen. Air flow running through the openings of the 3d printed cup suspended an 8-inch Styrofoam ball, allowing mice to run comfortably on it when head-fixed. Optical flow sensors (ADNS3080) placed below the 3d printed cup tracked the optic flow of the ball’s patterns through a small glass window at the cup’s base, which was then translated into ball movements via an Arduino Due. Light emitted from a DLP projector (Optoma) was reflected on a spherical mirror (Thorlabs) and then projected onto a custom-built Styrofoam toroidal screen, thus producing the VR environment scenes. The VR environment covered the visual field by ∼ 270⁰ horizontally and ∼80⁰ vertically. The VR environment was generated and controlled with ViRMEn^58^. Rewards were delivered by a miniature inert solenoid valve (Parker) and controlled with a NI-DAQ card (PCIe-6321, National Instruments).

### Behavior and training

Mice performed a task switch protocol^28^. Mice alternated in blocks between a navigation-based evidence accumulation task (‘towers task’)^45^ and a simpler task featuring the same virtual reality environment without requiring evidence accumulation (visually guided task). The virtual environment for both tasks was a 330-long T-maze comprising of 4 main regions/epochs, a 30-cm start region, a 200-cm cue region, a 100-cm delay region and the arm/reward zone at the end of the maze. The visual guide presented in the visually-guided task was always congruent with evidence (49 sessions, 9 animals). The statistics of the cues in the cue region were the same in both tasks. The towers positions were drawn from a Poisson distribution, such that the mean density for the cues on the rewarded side was 7.7 m^-1^ and for the distractors, i.e. the cues on the unrewarded side, was 2.3 m^-1^. These numbers were chosen such that the mean task difficulty was set to log(7.7/2.3) = 1.2^59^. The cues appeared when the mouse crossed the 10^th^ cm before the cues’ programmed location and disappeared 200 ms later. The maximum number of cues on each side was set to 16 and the minimum distance between two consecutive cues to 12 cm. On correct trials, a drop of 10% sucrose water was dispensed from a lick spout in multiples of 4 µL. In the towers task, the volume of the reward was typically 1.6 x 4 = 6.4 µL, whereas in the visually guided task was set to 1.2 x 4 = 4.8 µL. Occasionally on days of bad performance, animals were motivated to perform by increasing reward volumes to 2 or 3 times the initial amount. Even after reward adjustments, the difference in reward volumes between tasks was kept constant to an extra 1.6 µL for rewards in the towers. The end of a trial was followed by an intertrial interval after which mice were teleported back at the start of the maze. The intertrial interval was 3.5 s on correct trials, and 3.5 plus an extra 3 s, total ∼6.5 s, on error trials. Error trials were also signaled with a loud noise that played right after mice made a turn into the wrong arm. Each session started with the visually guided task as warm-up. Typically, warm-up lasted for ∼10 trials, unless performance was low, in which case the warm-up phase was extended until mice reached 85% correct within a rolling average of 40 trials. The length of subsequent task blocks varied for the towers between 40 and 60 trials and for the visually guided task between 15 and 30. Within a single session, mice typically performed ∼9-10 blocks and a total of 300-350 trials.

The training procedure was the same as previously described^45^. Mice were trained in the towers task with a shaping protocol comprising of 11 stages. Mice automatically progressed through the various training stages when certain criteria were met, such as recent performance from the last 40 trials, low bias and the quality of the animal’s trajectory. Mice reached the final stage within 4 to 6 weeks. As soon as mice were able to perform the towers task with a minimum performance of 70%, we switched them to the task switch protocol. Mice were able to switch between the towers and the visually guided task immediately, typically reaching block performance higher than 90% in the visually guided task and higher than 65% in the towers.

### Preprocessing

Preprocessing analysis was streamlined using DataJoint (DJ). Info about an imaging session (directories, date and time, tiff headers, other imaging parameters), synching between the neural data and behavior, imaging data preprocessing parameters, and resampled data were all ingested into DJ and fetched from DJ for any subsequent analysis.

For image preprocessing, we parsed the Tiff files into Tiffs for individual areas. Next, we used Suite2p on each area separately for image registration and ROI segmentation^60^. Only rigid registration was performed, which was sufficient to correct movement of 30 μm or less in x and y. Sparse mode was set to false, ROI diameter was set to 10 μm and maximum overlap between two ROIs to 0.2. We used a semi-automated approach to select for putative neurons with the Suite2p GUI. We used ‘skewness’ as a measure for positive transients and included neurons with at least a couple of transients throughout the session. We also used ‘compactness’ as a measure for neuron -disk-shapes, considering values smaller than 1.3. We also visually inspected many neuron traces using the registered movie. Most analyses were performed on all the neurons identified with the above criteria. The only exception was the sequencies, for which we used a similar selection criterion as in^17^. For these analyses we only included cells with at least 0.1 positive transients per trial: 1,899 in M2 (24% of all cells), 4,149 in RSC (40%), 2,635 in AM (30%). A transient was defined as the portion of the 𝛥*F*/*F* trace with magnitude at least 2 standard deviations above noise level and for the duration of at least 4 frames (∼300 ms). The noise level was defined as the FWHM of the neuron’s 𝛥*F*/*F* distribution.

Since we used online motion correction, fluorescence traces were overall stable. Therefore, we calculated 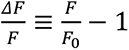, where *F_0_* is the modal value of fluorescence obtained from the whole session.

Given the block structure of our behavioral protocol, we assessed whether a session would be used for further analysis based on the behavioral performance on a block-by-block basis. To select ‘good’ blocks we used the following criteria: a) mean block performance above 60% for the towers task and 80% for the visually guided task, b) mean side bias less than 40%, defined as the absolute difference between percent correct in right versus left choices. In addition, time-out trials or trials in which the animal’s trajectory was longer by more than 10% of the nominal maze length were also excluded from further analysis. A block was deemed ‘good’ if it contained at least 3 trials passing the above criteria. Finally, we only analyzed sessions with at least 30 correct trials in each task. 46 out of an initial pool of 49 imaging sessions from 9 animals passed these criteria.

Synchronization between the neural and behavioral data was performed with the ScanImage I2C protocol. At each VR iteration, the VR computer sent a timestamp of an incoming packet, which the SI saved into the Tiff header of the current image frame. The incoming packet also contained information on the current acquisition, trial and virmen iteration. Since the VR frame rate was approximately 3 times higher than the imaging framerate, typically 3 I2C timestamps were saved in each imaging frame. To align behavior to the neural data, only the first VR timestamp was used. Ignoring additional VR timestamps for a given imaging frame led to an imprecision in the timing of behavioral events of the order of a few 10s of milliseconds, a timescale much smaller than the timescale of the Calcium indicator, and hence we did not attempt to correct it.

All ΔF/F traces and behavioral motor variables, such as running speed and view angle, were resampled as a function of a task-epoch axis. Each trial featured 5 main epochs: start, cues, delay, arm and intertrial interval (ITI). We determined the total number of bins per epoch based on its duration, such that the bin width was approximately the same across all epochs, between 150 ms and 200 ms. The total number of bins per epoch was: 4 bins at the start, 19 in the cues, 12 in the delay, 4 in the arm, 19 for the ITI, and another 18 for the extra ITI in error trials. Typically, most variables were resampled by taking the average within each bin. Only exception was the number of ‘tower’ cues, in which case we took the sum of all towers within a given bin. Subsequent analysis was performed on the resampled ΔF/F traces, and on correct trials only, unless otherwise stated (see below).

Manifolds were inferred from the ΔF/F time-series of correct and equalized trials between tasks. ΔF/F raw traces were also smoothed using an 11-frame Gaussian window (∼800 ms) and thresholded to 2σ, with σ estimated from each neuron’s entire time series^21^. The data were further preprocessed using PCA, keeping a total number of PCs that explained 95% of the variance.

### Behavioral analysis

Evidence was defined as the difference between number of towers presented on the right minus number of towers presented on the left (Δ(#R-#L)). Percent correct performance was defined as the number of choices/turns towards the side with most towers over the total number of trials. Side bias was defined as the difference in percent correct right trials versus percent correct left trials.

Modulation of behavior by evidence in each task was quantified by taking the difference in performance between trials with high evidence (Δ(#R-#L) > 5) versus trials with low evidence (Δ(#R-#L) ≤ 5). Variability across subjects was quantified by estimating the percentiles of that difference from the distribution across sessions and comparing them between tasks. Behavior was deemed more different between tasks if the difference between the 25^th^ percentile in the towers and the 75^th^ in the visually guided task was larger.Psychometric curves were estimated as previously described in ^45^. A sigmoid with 4 parameters was fitted to the right choices as a function of the binned evidence (#R - #L). Bin width was set to 3 and its mean value was weighed by the number of trials.

We compared the motor behavior between tasks by analyzing 3 key motor variables: the speed, view angle and turn onset, as previously defined in ^28^. Running speed was derived from the x-y displacement and the average run speed from averaging across positions in the maze stem (0 <y < 300 cm). Deviations in view angle (view angle SD) were quantified with the standard deviation across angles in the maze stem. The turn onset was estimated as the point when the view angle derivative was 3 standard deviations larger than baseline, where baseline was the mean view angle derivative between the start of the maze and the start of the delay (y = 200 cm).

### Cue-locked activity

This analysis was performed separately in each task and only on neurons with at least 0.1 positive transients/trial during the cue epoch. Each neuron’s ΔF/F trace was cross-correlated with the cues on the left or the right, during left or right correct trials respectively, in the interval [-1.5 1.5] seconds relative to the cues’ onset. To select neurons that were significantly locked to the cues, we estimated the cross-correlations with a shuffled dataset, where the position of cues was shuffled 100 times, while preserving their total number on each side. The significance criterion was defined as the z-scored difference between the cross-correlation peak from the actual trace versus the peak from the shuffled traces. Significance threshold was set to 2.

### Choice-selective sequences

Choice-selective sequences were obtained from correct trials only, and from neurons with at least 0.1 positive transients/trial. To compare choice-selective sequences between tasks we used a cross-validated approach. We used the towers task odd trials only to determine each neuron’s preferred side and position in the maze to fire. To determine each neuron’s preferred side to fire, we first located its firing field’s FWHM from the odd trials. Provided the FWHM was at least ∼1.5 s long, we performed a pairwise statistical comparison (paired *t*-test) at FWHM between left and right choice trials. A neuron was deemed left or right choice selective if its mean firing field during left or right trials was significantly different than the opposite side. For choice-selective neurons, their preferred position to fire was set to the location of peak activity on their preferred side. Whereas for cells that were active but not selective, their preferred position to fire was determined from both sides/choices. For each neuronal subpopulation, left-preferring, right-preferring or non-preferring, mean firing fields from the towers even trials or from all the visually guided trials were used to plot choice-selective sequences based on the order determined from the towers odd trials. Hence, the same ordering was maintained across tasks. In addition, all plotted firing fields were normalized by the same value, the maximum of the firing field obtained from the towers odd trials. The sequential activity across the population was captured with the ‘normalized rank in sequence’, that is the percentage of neurons across all choice groups with preferred position equal or smaller than a given position in the maze. . The firing field width at a given position in the maze was estimated from the test set only, by averaging the full width at half the maximum response (FWHM) across neurons preferring to fire at that position.

### Pairwise correlations

Pairwise correlations were performed on the single-trial, resampled ΔF/F traces from correct and equalized trials between tasks. Before estimating correlation coefficients, we first sorted neurons according to their preferred position to fire. We then concatenated neural traces from all 3 areas in the order M2-RSC-AM and estimated the pairwise correlation of each neuron with all other neurons for each task separately. The resulting correlation matrix is square and symmetric, containing the square and symmetric submatrices of the within-area pairwise correlations along the diagonal, and the rectangular across-area submatrices in the off diagonal. For the scatterplot of pairwise correlation coefficients in one task versus the other, we considered all possible pairs without repetition. R^2^ values for each session were obtained by linearly regressing the towers pairwise correlation coefficients from the visually guided coefficients. Average across and within correlations were estimated by taking the average across all elements of the corresponding matrices, except from the diagonal elements of 1s in the within pairwise correlation matrix.

An essential control when comparing pairwise correlations between two conditions is that any difference in pairwise correlations is not due to differences in the firing rates. We estimated each neuron’s mean firing rate in the towers or the visually guided task and compared those across the population on a session-by-session basis.

### Linear task subspaces

For each task, we determined the directions and amount of maximum variance explained by estimating the eigenvectors of each task’s pairwise correlation matrix that corresponded to the largest eigenvalues. Since the pairwise correlations can be thought of as the normalized covariance matrix, this process is the same as performing principal component analysis (PCA) on the standardized data matrix with rows #trials X t and columns # neurons from all 3 areas. We performed the eigendecomposition using a cross-validated approach, which would then allow us to make a fair comparison between the variance captured within-task versus across-tasks. For each task, eigenvectors/principal components (PCs) were obtained from the odd trials only. Then the variance captured was determined by projecting the held-out data from the even trials onto the ‘odd-trial’ eigenvectors. In matrix notation: 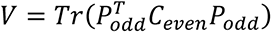 is the matrix trace, 𝐶 is the covariance obtained from even trials and𝑃 are the eigenvectors obtained from odd trials. We looked at the projections from the first 20 PCs which captured approximately 20% of the variance. To assess the difference between projections onto one task versus the other, we defined the variance explained (VE) index as: 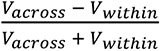, that is the normalized difference of the variance captured when projecting onto the same basis either data from the other task (*𝑉_𝑎cross_*) or data from the same task (*𝑉_within_*).

For the comparison across areas, we equalized the total number of cells in each area, by randomly subsampling without replacement 10 times. The VE indices reported were obtained by averaging across subsampling iterations. Since the within-area pairwise correlation matrices are also square and symmetric we performed the same analysis as above for each area individually. Approximately half the number of PCs was required to capture a similar percentage of variance (20%) compared to all the data concatenated across areas. Instead to estimate the variance captured by the across-area PCs we performed SVD and estimated the variance captured either by the left or the right-singular vectors. Approximately a 3^rd^ of the total number of PCs was required to capture a similar percentage of variance (20%) compared to all the data.

### Nonlinear manifold inference

To infer low-dimensional latent dynamics from the population neural activity we used MIND (‘manifold inference from neural dynamics’^50^). Briefly, MIND recursively partitions the neural state space into local neighborhoods using random forests^61^. Within each sufficiently small local neighborhood, MIND learns a distribution over successor states given the current state using probabilistic PCA^62^. As with any random forest algorithm, the learned distributions from a large set of randomized trees are combined, giving rise to an ensemble model of transition probabilities from a given neural population state to the next. These transition probabilities are then used to define local distances, such that the distance assigned between two states is smaller if the probability of directly transitioning between these two states is higher. The global distance was then estimated across all possible pairs of states from the local distances: given a non-zero transition probability from state i to state j, then states i, j are vertices on a directed graph G with global distance the symmetrized geodesic distance on G. d-dimensional intrinsic dimensions/latents were obtained using a form of metric multidimensional scaling under the constraint that distances between points on the manifold are approximately preserved. Forward mapping from the latents to neural activity and its inverse was performed with local linear embedding (lle).

Similar to Nieh et al., the orientation that best split the data in each node was selected from a total of 20 possible orientations and the minimum number of leaves per random tree was set to 500. We used the ‘landmark’ approach in ^50^: after transition probabilities have been estimated on the full dataset, the manifold structure is learned from a ‘subset of landmarks’, whose size was set to 1/7^th^ of the full size. Best values for the lle hyperparameters, the regularization strength and number of local neighbors, were searched within a given range using 10-fold cross-validation. For each dataset, the manifold was embedded into d dimensions in the range 2 to 8. To assess how well a given set of latents could predict the activity of single neurons we used a cross-validated approach. We used 85% of the trials to fit the manifold, and the remaining 15% to embed the manifold into d = 2-8 dimensions. The d latents were then mapped into the full neural state space of dimension n >> d, giving rise to n reconstructed neural activity traces. The reconstructed traces were preprocessed the same way as the actual data (see preprocessing). The reconstruction score was defined as the Pearson’s correlation coefficient between the concatenated actual data and the reconstructed traces.

In the first half of the manifolds analysis (Figure 2) we applied MIND on the whole session, thus estimated one lower-dimensional subspace for both tasks and the ITI interval. In the second half (Figure 4) we treated the two tasks independently, obtaining one manifold for each task. Because the two manifolds were used to decode behavioral variables such as evidence and position during the cues and delay epochs (see decoding from aligned task manifolds), manifold extraction was restricted to periods when the maze was on^21^.

In 2 out of 46 sessions we experienced convergence issues, and therefore the two sessions were excluded from further analysis with MIND.

### Pairwise firing field overlap on the manifold

‘Manifold firing fields’ were defined as the activity of each neuron expressed as a function of the manifold latents obtained from both tasks. For each neuron, we first estimated the center of mass (CoM) of its manifold firing field, 𝐶_𝑘_, and the second moment, 𝐼_𝑘𝑘_, that is the spread of points around 𝐶_𝑘_ weighted by that neuron’s ΔF/F. Next, for each pair of neurons (𝑗, 𝑘), we estimated the second moment of neuron 𝑗 relative to the CoM of neuron 𝑘, 𝐶_𝑘_, 𝐼_𝑗𝑘_ ≡ 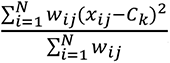, where *𝑥_i𝑗_* is the i’th sample of the j’th neuron’s firing field manifold coordinate and 𝑤_𝑤*i*𝑗_ is the i’th sample of the j’th neuron’s ΔF/F. Finally, we defined the firing field overlap ratio for the pair (𝑗, 𝑘) as 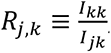, so that a ratio of 1 means perfect overlap, a ratio less than 1 means partial overlap whereas a ratio larger than 1 means that the firing field of neuron 𝑗 is contained within the firing field of neuron 𝑘. Note that 𝑅_𝑗,𝑘_ ≠ 𝑅_𝑘,𝑗_ and therefore for a given pair we are reporting the mean: (𝑅_𝑗,𝑘_ + 𝑅_𝑘,𝑗_)/2. To obtain histograms of the difference in firing field overlap between tasks, we estimated the mean overlap of each cell with all other cells and then averaged across manifold dimensions. Finally, to obtain summaries across sessions (n = 44), in each session we estimated the grand mean across all pairs of neurons and latent dimensions (n = 5).

### Cross-validated evidence sequences

Evidence sequences were obtained from correct and equalized number of trials between tasks, and from neurons with at least 0.1 transients/trial. Firing fields in the evidence domain were estimated by binning ΔF/F in 25 evidence bins of bin size 1 in the interval [-12, 12]. For each neuron and task we estimated a pair of mean firing fields, one obtained from odd and one from even trials. Both mean firing fields were smoothed using a Gaussian window with a length of 5 bins and a σ of 1 bin^21^. Mean firing fields from even trials, the test set, were used to plot the sequences based on the order determined from odd trials, the train set. In addition, firing fields from both odd and even trials were normalized by the same value, the maximum of the firing field obtained from odd trials. We selected cells whose firing fields between odd and even trials were positively correlated with a p value < 0.01 either in the towers or the visually guided task. The firing field width of the two subsets of cells was estimated from even trials only, by counting the number of evidence bins at half the maximum response (FWHM), separately for the towers or the visually guided task.

### Evidence decoding from the latents

To obtain an evidence axis within the low-dimensional latent space, we decoded evidence linearly from odd-trial latents and projected the corresponding even-trial latents onto it. Decoding was performed separately for each task. We then estimated firing-field overlap along these projections, concentrating on the subpopulation of neurons selected in the evidence sequences analysis.

### Decoding from aligned task manifolds

We assessed the similarity in the geometric structure between task manifolds using the approach previously described in ^21^. We considered the case of n = 5 dimensions, the minimum number of dimensions with reconstruction scores close to the maximum across tasks and areas. In this five-dimensional space each manifold forms a special orthogonal group (SO(5)) with 10 free parameters which can vary while preserving its structure. The 10 free parameters correspond to the 10 possible rotations in the 5-dimensional space. A given task manifold was deemed optimally aligned to the second task manifold if it gave the best prediction based on a decoder trained on the second manifold. In technical terms, we trained two linear decoders, one for position and one for evidence, from the latents of one task using even trials only. We then tested each decoder on the held-out portion of the same task’s latents (the odd trials) and obtained an estimate for the decoding accuracy within task (Pearson correlation). Next, we used the even trials of the second manifold’s latents to determine the rotation transformation that maximized the decoding accuracy for both evidence and position. And finally, we tested the predictions of the evidence and position decoders on the second task’s rotated latents obtained from the odd trials, which yielded the decoding accuracy across tasks. We tested the alignment and decoding either by aligning the towers task to the visually guided manifold, or vice versa.

### Density scatterplots

A grid of 200 x 200 bins in the range [min(X), max(X)] and [min(Y), max(Y)] was used to count the number of elements within each bin. The grid was then smoothed in 2d with a smoothing parameter λ = 20 and normalized by its maximum value. The smoothed normalized values determined the color of each pair of points (X_i_, Y_i_).

### General statistics

Distributions were summarized either with their mean ± S.E.M., or the median ± S.E.M. In the latter case, S.E.M. is defined as the difference between the 84^th^ and the 16^th^ quantile divided by √𝑛, with 𝑛 being the total number of samples. For all tests comparing the mean or median of distributions across sessions, we used mixed effects models, with the variable of interest being the fixed effect predictor and subjects being the grouping variable. Since we recorded multiple sessions per subject, the random effect was modeled as a sparse matrix with 1’s indicating which sessions belonged to a given subject.

The binomial confidence interval reported in the psychometric data was obtained from Jeffrey’s method ^45^.

Bootstrapping was performed by randomly sampling with replacement 1,000 times. P values were defined as the fraction of bootstrap iterations with slope ≥ 0.

## Acknowledgements

This work was supported by the EMBO postdoctoral fellowship ALTF 384-2020 (to E.M.D.), the US National Institutes of Health (NIH) grant U19NS132720 (to D.W.T. and C.D.B.), the Simons Foundation grant SCGB AWD543027 (to D.W.D.), the NIH grants K99MH120047, R00K99MH120047 (to L.P.), and the CV Starr fellowship and Burroughs Welcome CASI award (to M.S.). We thank Marius Pachitariu for help with the implementation of online motion correction, Meeanakshi Ashokan, Flora Bouchacourt and Mark Ioffe for thoughtful comments on the manuscript, P. Dylan Rich and all members of the Tank lab and the U19 collaboration at Princeton for valuable feedback on the study.

## Aut hor cont ributions

E.M.D. L.P. and D.W.T. conceived the project, E.M.D. collected the data, E.M.D. L.P. and A.A.R. performed pilot experiments, E.M.D. performed the analysis with input from L.P., M.S., C.D.B. and D.W.T, S.Y.T. designed the optical setup, E.M.D. wrote the manuscript with input from L.P., M.S., A.A.R, C.D.B. and D.W.T. D.W.T. supervised the work.

## Dat a and code availabilit y

The full dataset and source code will be publicly released upon publication of this work

---

**Supplementary Figure 1.**
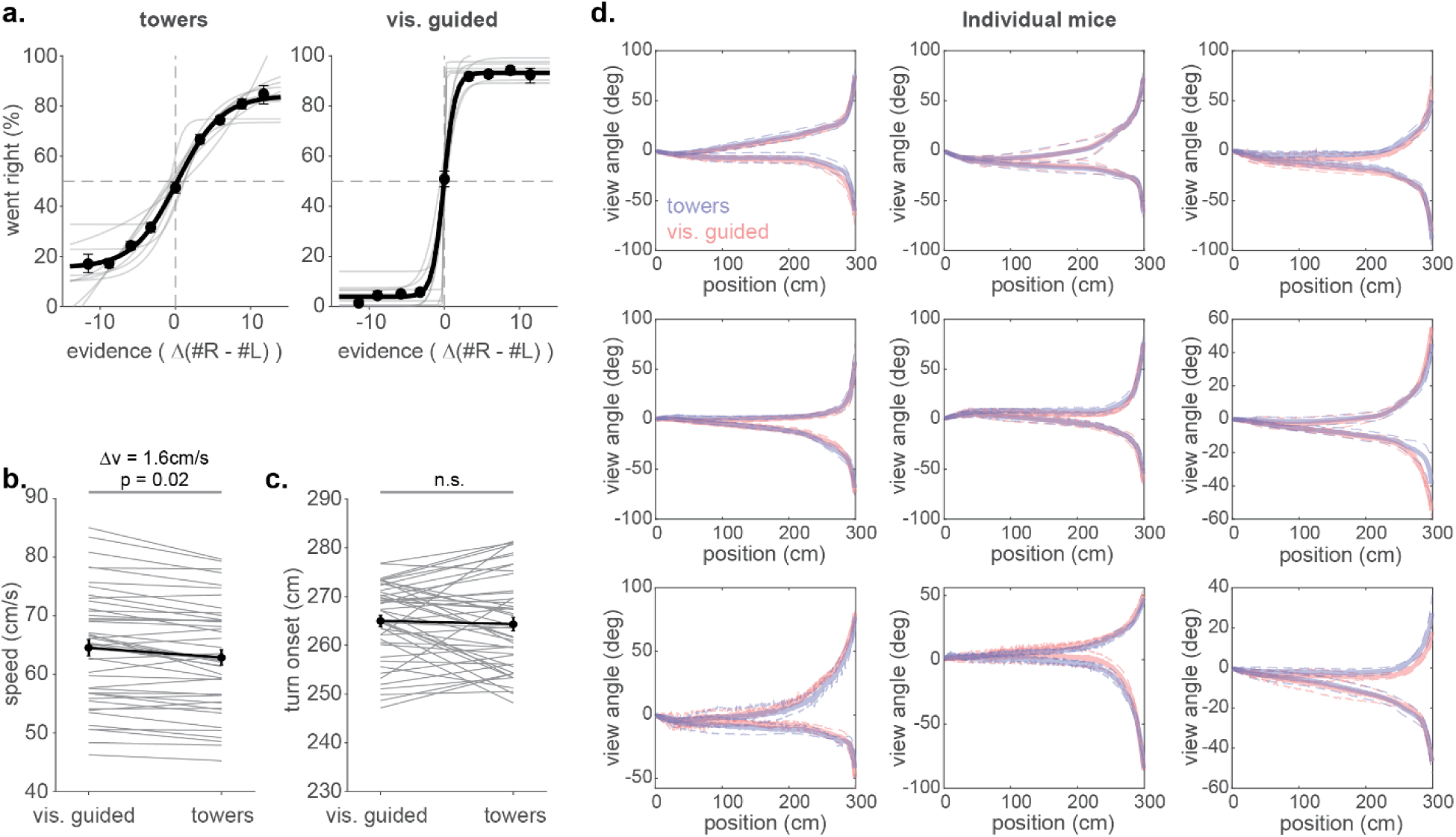
(a) Psychometric curves in the towers task (left) or the visually guided task (right) across mice. Gray lines are from each mouse (n = 9). Black datapoints are from data pooled across all mice, black line is the fit to these datapoints. Error bars: binomial confidence interval. **(b)** Paired comparison of the mean speed between tasks within the same session (n = 46; gray lines). Black line and datapoint: mean across sessions; error bars: S.E.M; p value from mixed effect model **(c)** Same as in **(b)** for the turn onset. **(d)** Each mouse’s view angle as a function of position in the maze during the towers task (blue) versus the visually guided task (red). Solid lines: mean across sessions; dashed lines: S.E.M.

**Supplementary Figure 2.**
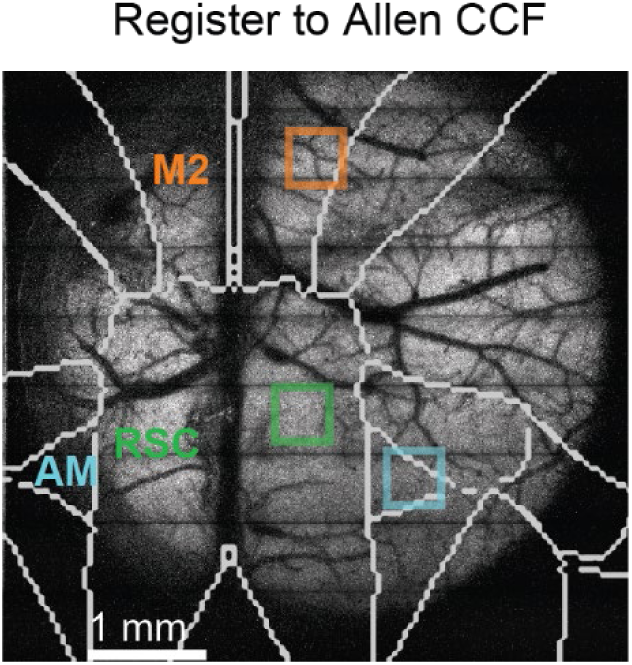
Same field of view as in Figure 1b with registered Allen CCF contours superimposed (white), and ROIs from individual areas, M2 (orange), RSC (green) and higher visual AM (cyan), targeted over the right hemisphere. The names of individual areas are also shown over the left hemisphere.

**Supplementary Figure 3.**
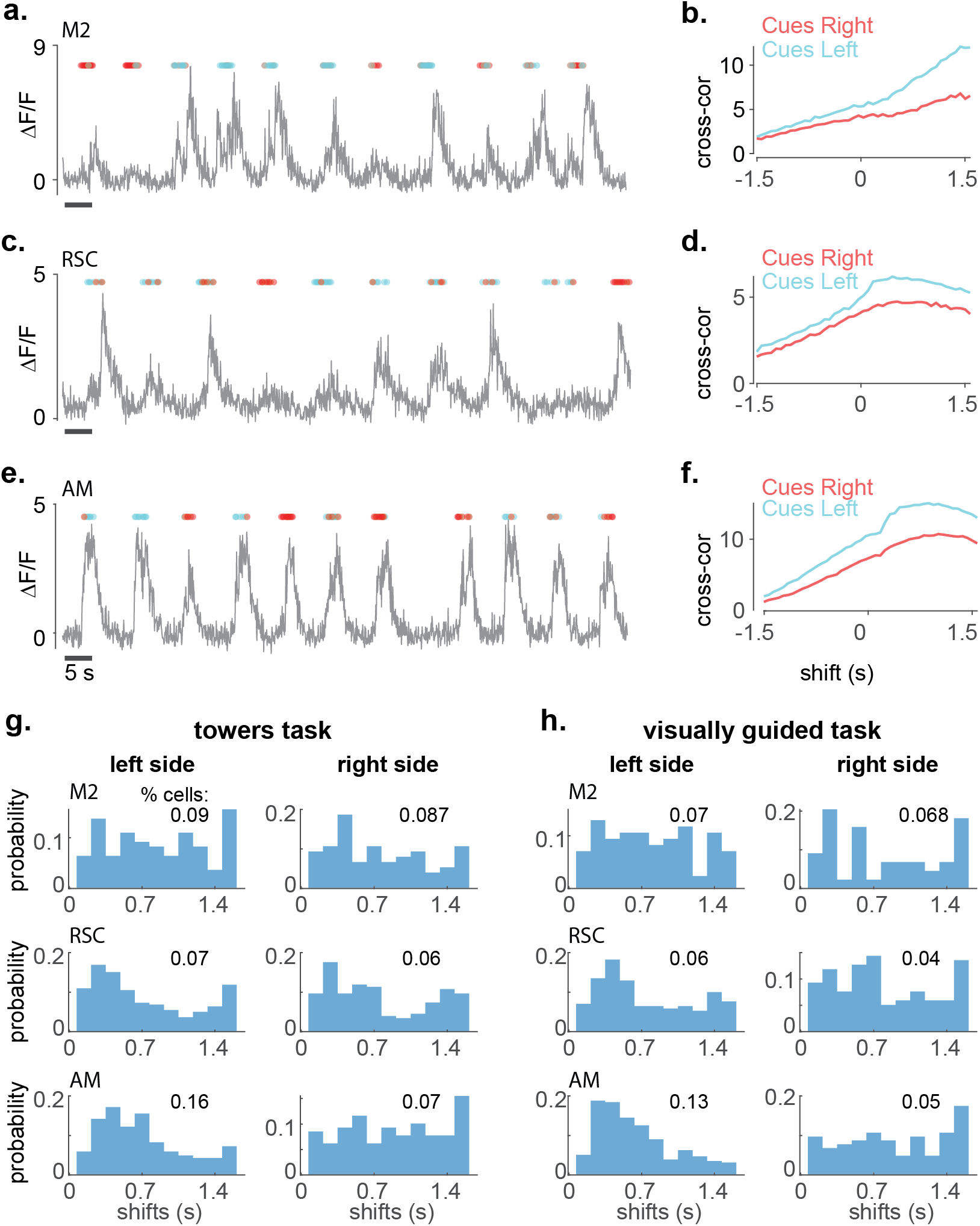
(a) example ΔF/F time series of a cue locked neuron from area M2 (circles: onset of left (cyan) or right (red) cues). **(b)** Cross-correlation of the same neuron’s ΔF/F with the left cues during correct left trials (cyan), or with the right cues during correct right trials (red). **(c) – (d)** Same as **(a) – (b)** for RSC. **(e) – (f)** Same as in **(a) – (b)** for AM. **(g)** Normalized distributions of the time point relative to the cues onset at which each neuron’s cross-correlation peaks (‘shifts’), in area M2 (top), RSC (middle) and AM (bottom). Distributions were estimated separately for correct left or right trials during the towers task. **(h)** Same as in **(g)** for the visually guided task.

**Supplementary Figure 4.**
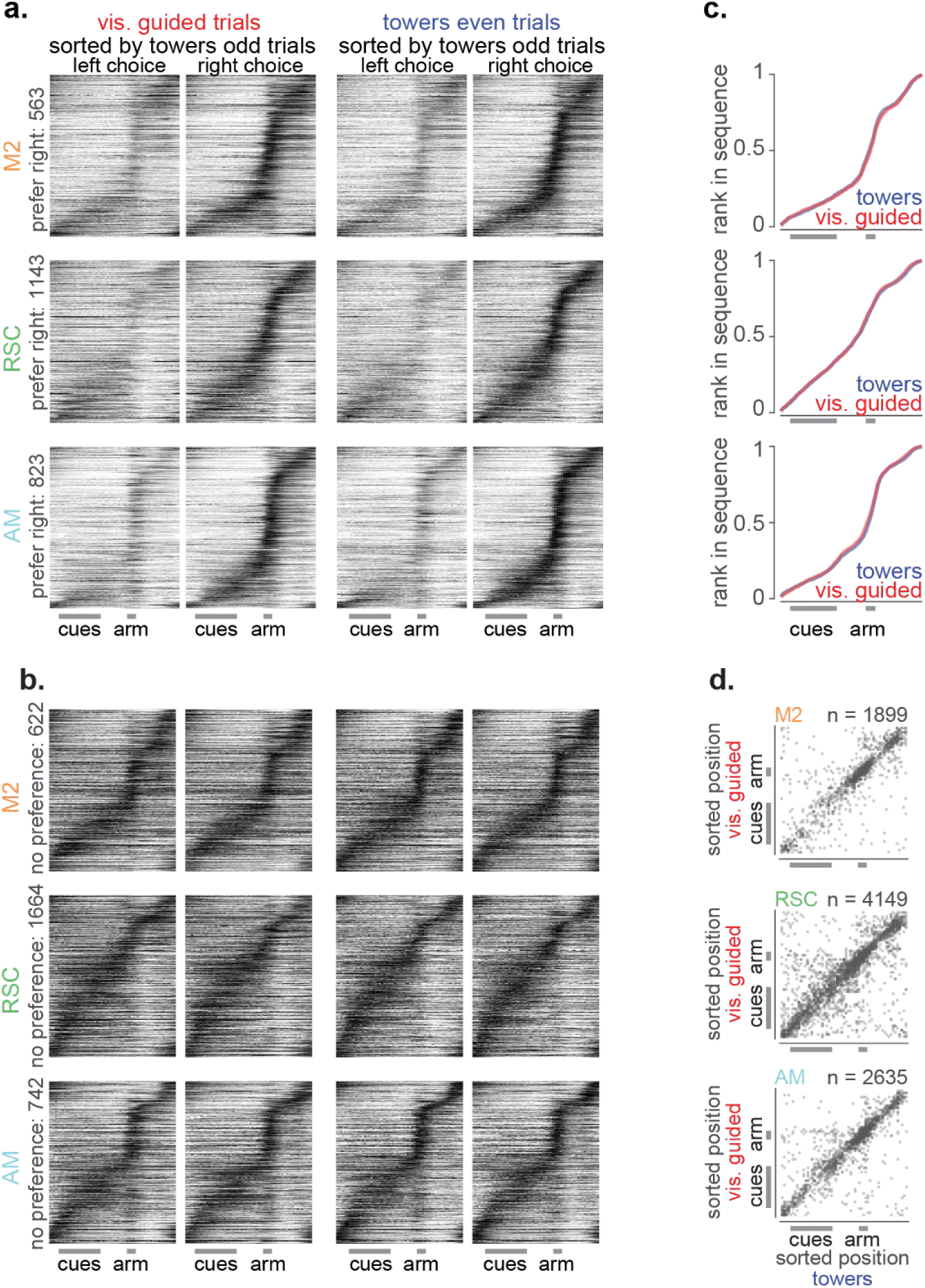
(a) Cross-validated right choice-selective sequencies in the visually guided task (left two columns) and the towers task (right two columns) in premotor M2 (top), RSC (middle) and AM (bottom). Neurons in both tasks were sorted based on the order obtained from the towers odd trials (‘train’ set). The plotted mean firing fields were obtained from all trials in the visually guided task and from the even-numbered trials in the towers task (‘test’ set). Sequencies span the whole trial plus the intertrial interval. x-axis gray scale bars: cues, arm. **(b)** Same as in **(a)** for neurons with no preference for left or right choice. **(c)** Percent of neurons with preferred position equal or smaller than a given position in the maze obtained from the test sets of each task (top: M2; middle: RSC; bottom: AM; blue: towers task, red: visually guided task). **(d)** The neurons’ preferred position to fire during the towers task versus the visually guided task obtained from the test (even) trials and pooled across choice selectivity categories.

**Supplementary Figure 5.**
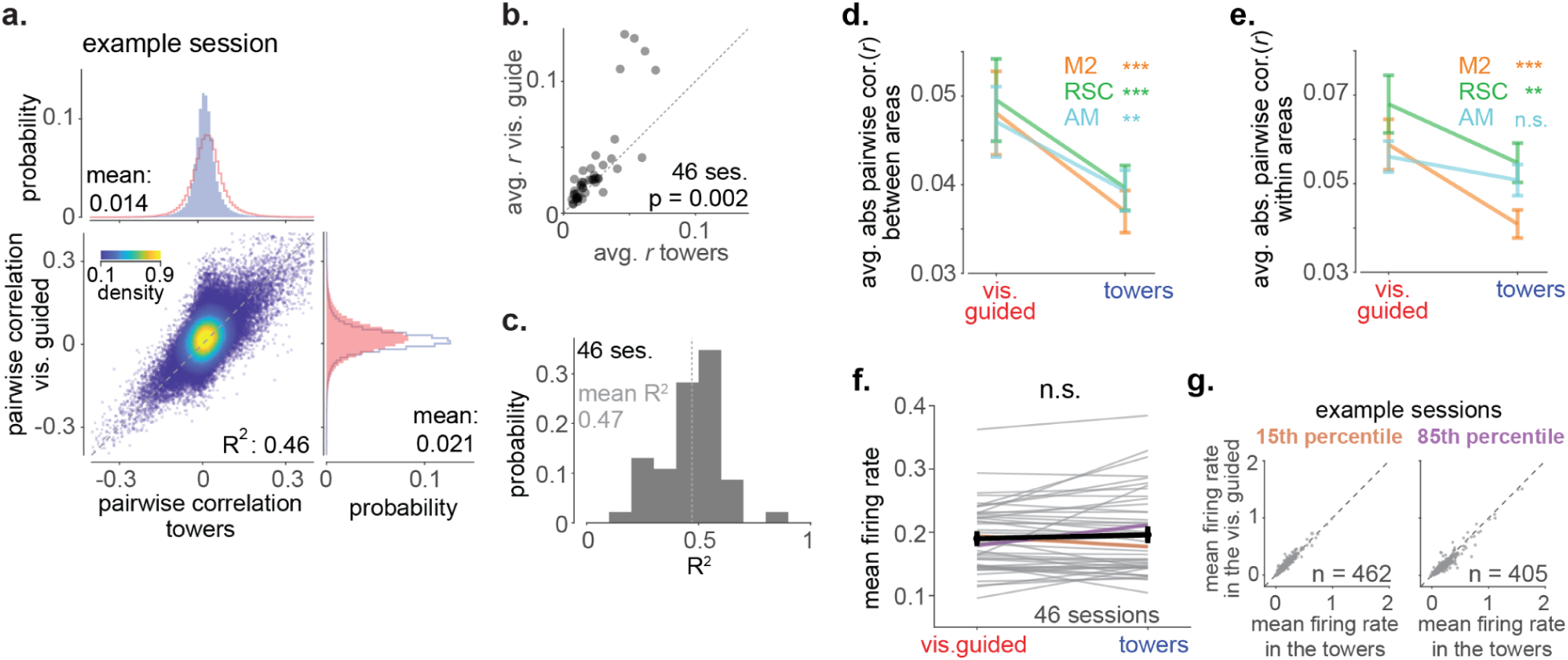
(a) Density scatterplot of the pairwise correlations in the towers against the pairwise correlations in the visually guided task. Data points correspond to all matrix elements lying above the diagonal in Figure 1(g). The histogram on top shows the marginal pairwise correlation distribution for the towers task (mean = 0.014; light blue), with the distribution for the visually guided task superimposed as a stairstep plot (pink). The histogram on the right shows the marginal pairwise correlation distribution for the visually guided task (mean = 0.021; pink), with the distribution for the towers task superimposed as a stairstep plot (light blue). **(b)** Average pairwise correlations in the towers versus the visually guided task. Each datapoint corresponds to one session. (n = 46, p = 0.002 from mixed effects model). **(c)** R-squared (R^2^) distribution quantifying the linear relationship between correlations in the towers versus the visually guided task across sessions (n = 46, mean: 0.47). **(d)** Average across-area absolute pairwise correlations in the towers versus the visually guided task for each area (p values from mixed effects model, M2: p = 0.0002, RSC: p = 0.0009, AM: p = 0.0035; error bars indicate S.E.M.) **(e)** Average within-area absolute pairwise correlations in the towers versus the visually guided task for each area (p values from mixed effects model, M2: p < 10^-4^, RSC: p = 0.0096, AM: n.s.; error bars indicate S.E.M.). **(f)** Paired comparison of the mean firing rate between tasks within the same session (n = 46; orange line: session from the 15^th^ percentile of that distribution; purple line: session from the 85^th^ percentile; gray lines: all other sessions). Black line and datapoint: mean firing rate across sessions; error bars: S.E.M; p value from mixed effect model. **(g)** Mean firing rate in the towers task against the visually guided task across all neurons recorded in two example sessions. Sessions were selected based on the distribution of mean firing rates across sessions shown in **(f).** Left: 15^th^ percentile; Right: 85^th^ percentile.

**Supplementary Figure 6.**
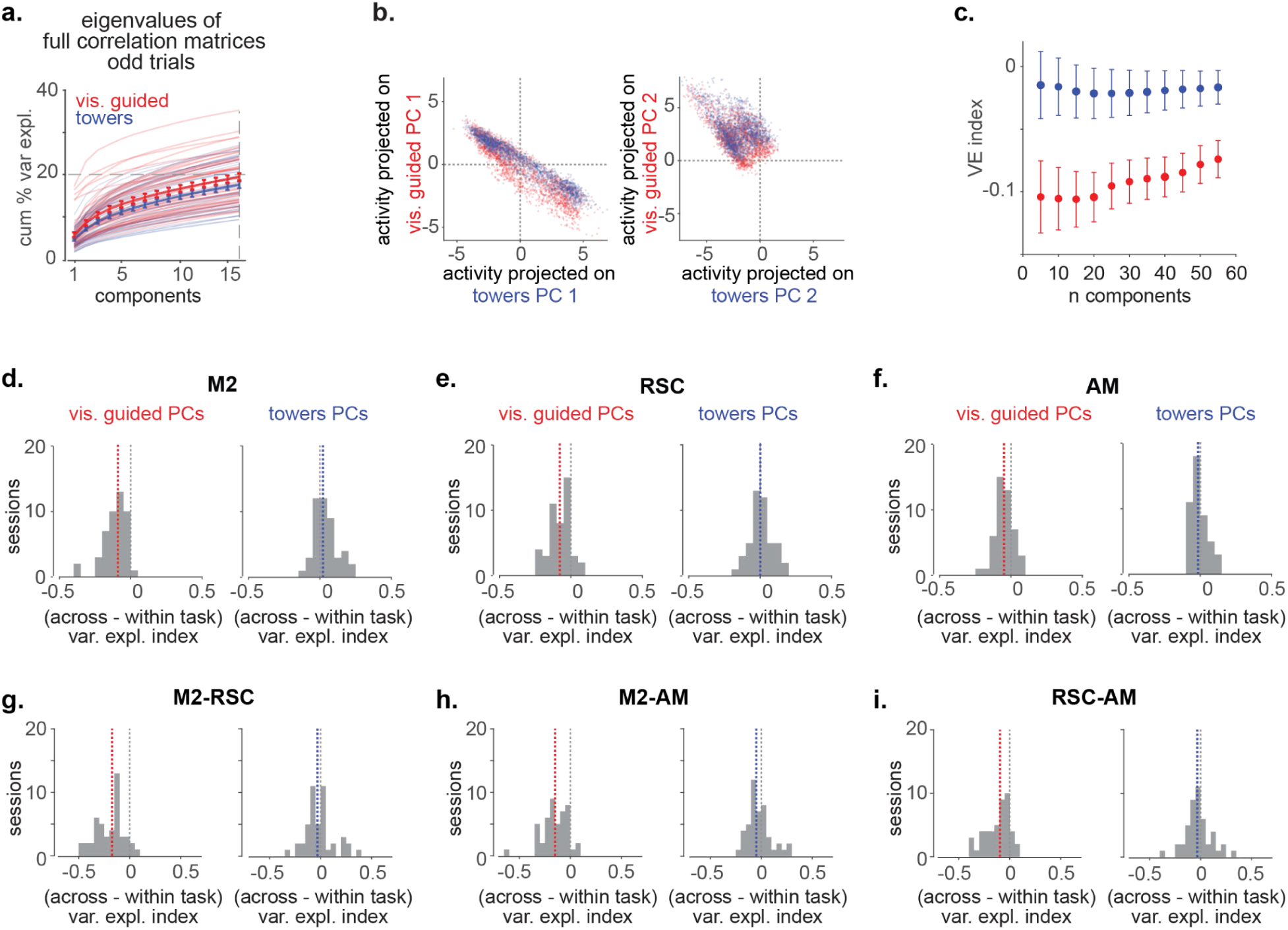
**(a)** Cumulative percentage of variance explained across the first 15 sorted eigenvectors of each task’s correlation matrix, obtained from the odd trials only (red thin lines: individual sessions during the visually guided task; blue thin lines: the same individual sessions but during the towers task; thick lines average across sessions for each task; error bars: S.E.M.; difference between tasks: 1^st^ component: p = 0.004, all other components: n.s., mixed effects model). **(b)** Projections of the even trial-to-trial neural activity in the towers (blue datapoints) or the visually guided task (red datapoints) onto the first (left) or the second (right) PC of the towers task plotted against the same PC of the visually guided task (same example session as in Figure 1**(g, j)**). **(c)** Variance explained (VE) index as a function of total number of components used to define eigenvectors/ PCs in each task (blue: towers task; red: visually guided task) **(d)** Distribution of the variance explained index across sessions and between tasks for M2 (left: visually guided, red dashed line: median; right: towers, blue dashed line: median). **(e) – (f)** Same as in **(c)** for RSC and AM. **(g) – (i)** Same as in **(c)** for the pairwise correlations between pairs of areas (M2-RSC, M2-AM, RSC - AM).

**Supplementary Figure 7.**
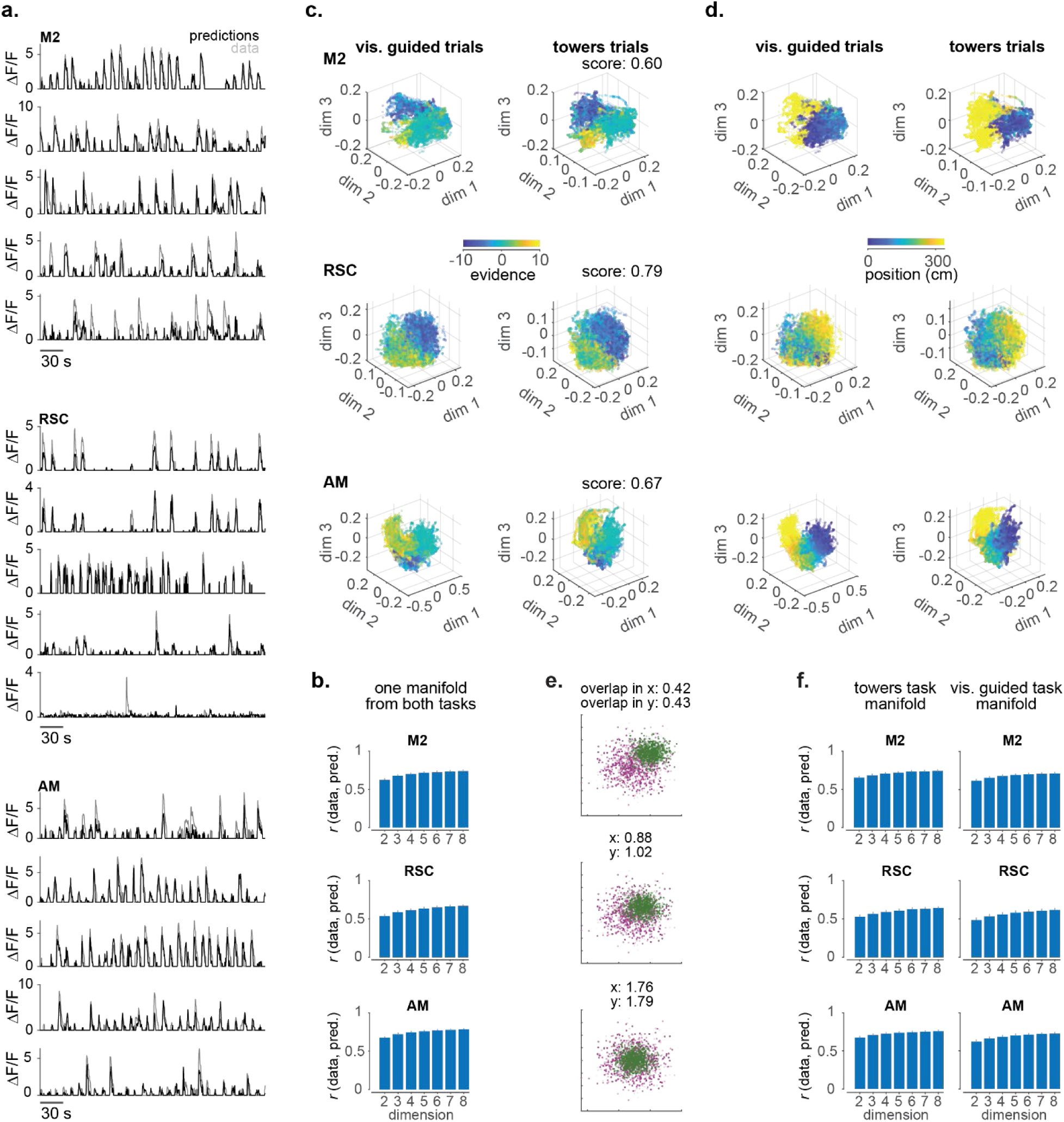
(a) Example traces and MIND predictions from the first 5 neurons in the dataset, in M2 (top), RSC (middle) and AM (bottom). Traces (gray) and MIND predictions (black) were smoothed using an 11-frame Gaussian window (∼800 ms) and thresholded to 2σ, with σ estimated from each neuron’s entire time series.**(b)** Average Pearson’s correlation, *r*, between single-neuron measured activity and the MIND predictions (‘reconstruction score’) as a function of total number of latents in M2 (top), RSC (middle) or AM (bottom). **(c)** Evidence (Δ(#L - #R)cues) plotted on the first three dimensions of the 5-dimensional embedding of a manifold obtained from each cortical area in both tasks. Submanifolds corresponding to matched number of correct trials in the visually guided task (left) and the towers task (right) were plotted separately. ‘score’ denotes the correlation coefficient between the reconstructed and the actual single-neuron traces (top: M2; middle: RSC; bottom: AM). **(d)** Same as in **(c)** for position. **(e)** Cartoon depicting the quantification of the overlap between two clouds of points. The datapoints from each cloud are drawn from a random distribution. The mean of the green cloud varies across panels, so that the two clouds partially overlap (top), fully overlap (middle) or the green cloud is contained within the purple cloud (bottom). x: overlap ratio along latent 1, y: overlap ratio along latent 2. **(f)** Same as in **(b)** after applying MIND separately on each task.

**Supplementary Figure 8.**
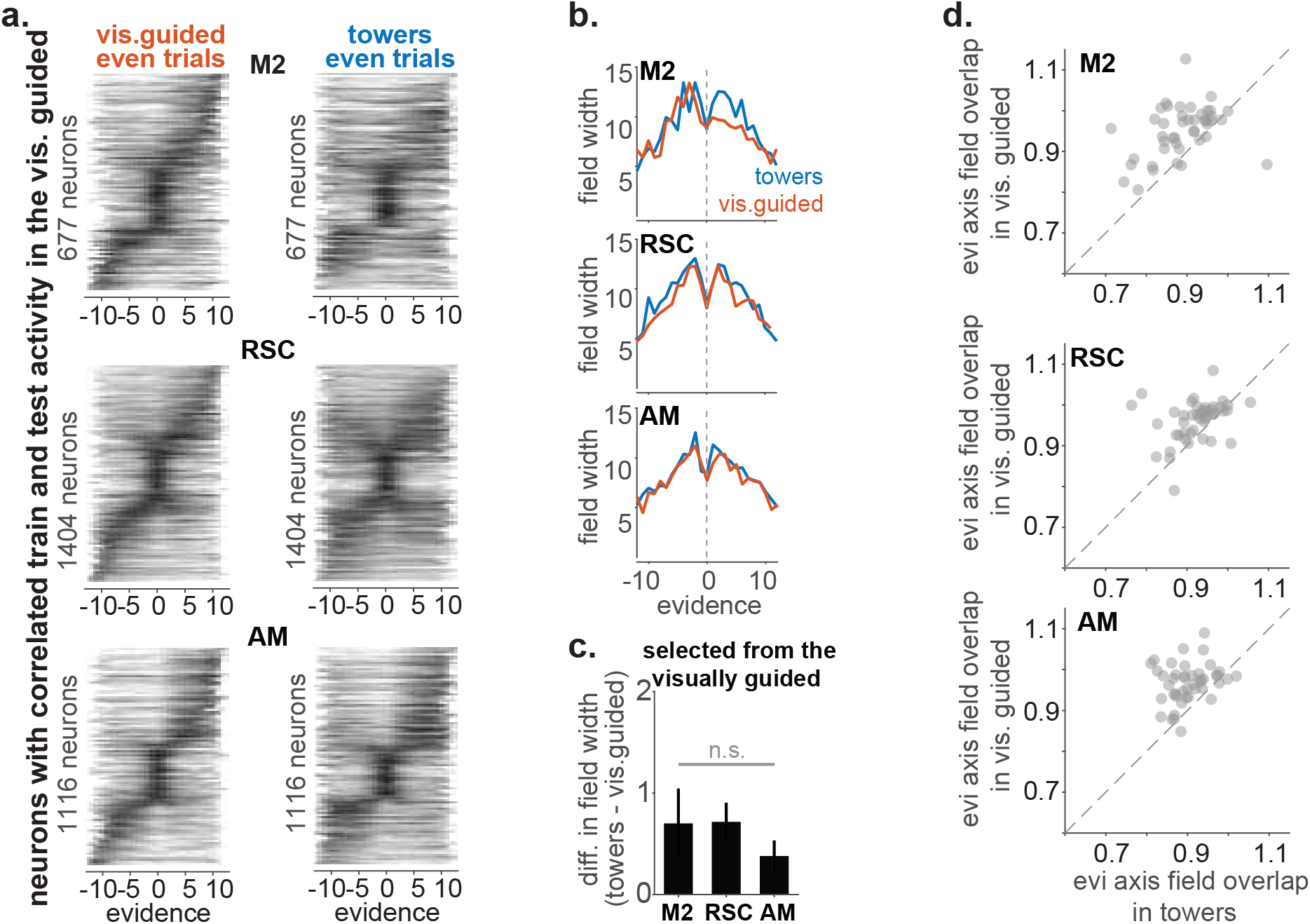
(a) Cross-validated sequences as a function of the evidence axis obtained from neurons with correlated train and test activity in the visually guided task (top: M2, middle: RSC, bottom: AM). Plotted sequences are from the even trials of the towers (right) or the visually guided task (left), while the ordering of neurons was determined from the odd trials of each task. **(b)** Mean field width of the neurons in **(a)** as a function of the evidence axis in the towers (blue) or the visually guided task (red). **(c)** Difference in field width in the towers versus the visually guided task averaged across all evidence values. **(d)** Mean manifold firing field overlap along the evidence axis, comparing the towers task to the visually guided, restricted to neurons forming sequences in the towers (same neurons as in Figure 3a). The evidence axis was obtained by linearly decoding evidence from the 5-dimensional latent space.

**Supplementary figure 9.**
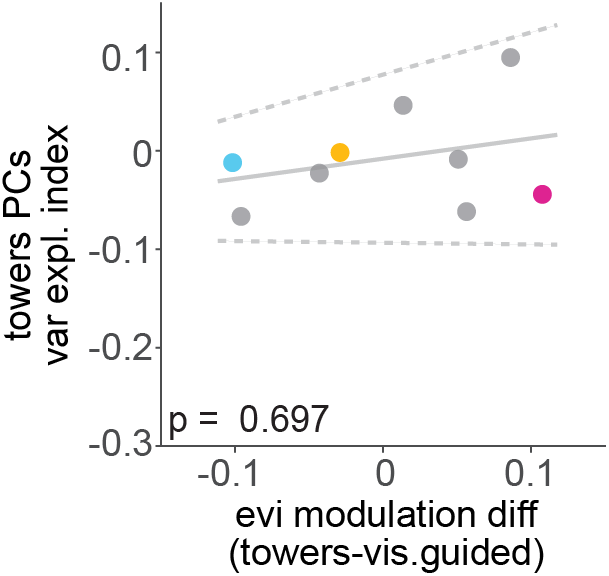
For each subject (n = 9), the evidence modulation difference (x axis) is plotted against the session-averaged variance explained index of the towers linear subspace. Unlike the linear visually guided subspace (Figure 5b), the magnitude of the variance explained index does not correlate with behavior. (solid line: fitted line averaged across bootstrapped iterations, dashed lines: ± 2 standard deviations). Colored datapoints: example mice shown in Figure 5a, gray datapoints: all other mice.

